# E2f coordinates the cell cycle and cell fate of hematopoietic progenitors to drive stress myelopoiesis

**DOI:** 10.1101/2025.06.16.659412

**Authors:** Annick Cachon, Gabriella Oliva, Eunsun Kim, Katharina E. Hayer, Yashvi Narravula, Lauren Hino, Stephanie Kjelstrom, Joanna Tober, Kai Tang, Nancy Speck, Patrick Viatour

## Abstract

During inflammation, cytokines such as Tnfα, Il1β and Il6 stimulate hematopoietic stem and progenitor cells (HSPCs) to proliferate and accelerate the production of inflammatory cells from the myeloid lineage through a process called stress myelopoiesis. Genetic inactivation of repressors of cell cycle activity in HSPCs is sufficient to recapitulate stress myelopoiesis, suggesting that the cell cycle activity and cell fate decision of proliferative HSPCs are coordinated by a cell intrinsic mechanism. However, the nature of this mechanism remains unknown. Here, we show that E2f simultaneously regulates proliferation of HSPCs, repression of alternative cell fates via the activation of *Suz12*/*Ezh2*-containing PRC2 complex and promotion of the myeloid fate by enhancing the signaling activity of CD131, the common βchain receptor for βcytokines (i.e. Il3 and Gm-Csf). Accordingly, dual inhibition of Ezh2 and βcytokine signaling activity in a preclinical model of colitis represses stress myelopoiesis and restores colon homeostasis. Our results suggest that dual targeting of βcytokine signaling and Ezh2 activity represents a novel therapeutic strategy to repress the production of pro-inflammatory myeloid cells in a wide spectrum of inflammatory diseases.

## INTRODUCTION

“Inflammatory diseases” (IDs) are pathologies of different etiologies characterized by increased expression of pro-inflammatory mediators and aberrant inflammatory cell activity. Pro-inflammatory mediators progressively generate systemic inflammatory conditions associated with significant co-morbidities that are poorly understood^1 6, 2^. This clinical evidence strongly suggests that chronic inflammatory conditions, while often initiating from one organ and clinically managed in a siloed manner, ultimately morph into multi-organ disorders requiring a comprehensive therapeutic approach. Current drug development strategy focuses on the inhibition of inflammatory cell activity. Drugs such as steroids, aminosalicylates and biologics have shown clinical benefits for ID patients. However, side effects and resistance limit their efficacy, urging for a paradigm shift to generate novel ID therapy^3^. In this perspective, pro-inflammatory myeloid cells are short lived, suggesting that inhibiting their production to starve the inflammatory process may represent a novel therapeutic strategy. Depletion of inflammatory myeloid subpopulations improves disease severity in pre-clinical inflammatory models^4–7^, providing a rationale for this strategy. However, the identity of the mechanism(s) driving the production of inflammatory myeloid cells remains elusive.

During homeostasis, highly quiescent long-term (LT)- and short term (ST)- hematopoietic stem cells (HSCs) occasionally enter cell cycle to self-renew or differentiate into lineage-primed multipotent progenitors (MPPs; megakaryocyte-primed MPP2s, granulocyte/monocyte-primed MPP3s and lymphoid-primed MPP4s ^18–21^), which further differentiate into lineage-restricted progenitors to ultimately generate terminally differentiated myeloid and lymphoid cells. Inflammatory conditions increase the expression of a common set of mediators (LPS, Tnfα, Il1, Il6, -Csf family, etc) that bind to receptors expressed at the surface of HSPCs to trigger stress myelopoiesis, which is characterized by sustained exit of HSPCs from quiescence, enhanced production of inflammatory myeloid cells and repression of alternative hematopoietic lineages.

Rb family proteins (Rb, p107 and p130) maintain cellular quiescence by repressing the transcriptional activity of the E2f family members (E2f1-8). Beyond its canonical role in cell cycle activation ^8–12^ ^13–18^, studies suggest that E2f also has non-canonical transcriptional functions during cell division^19,20^; however how they are integrated with its canonical cell cycle activity remains poorly characterized. Genetic inactivation of repressors of proliferation (*Rb* family, *p21*^21^, *p27*/*p57*^22,23^, *Pten*^24^, *Mek1*^25^, *Foxo family*^26^, etc) or overexpression of *Cyclind1*/*Cdk4* complex^25^ is sufficient to trigger the sustained HSPC proliferation and rapid myeloid expansion at the expense of other blood lineages that characterize stress myelopoiesis. This evidence suggests that derepressing E2f activity is sufficient to activate a cell intrinsic mechanism (i.e independent of the nature of the extrinsic stimuli that disrupts the Rb/E2f interaction) that drives stress myelopoiesis. However, the nature of this cell intrinsic mechanism remains unknown, which represents a major gap in our understanding of stress myelopoiesis and impedes our translational efforts to target inflammatory cell production for ID therapy.

Using a mouse model that enables the conditional inactivation of the *Rb* family, we show that E2f coordinates proliferation, the promotion of myelopoiesis and the repression of alternative hematopoietic cell fates via the simultaneous activation of its canonical cell cycle program as well as the expression of *Csf2rb* (which codes for CD131, the common βchain receptor for Il3&Gm-Csf) and *Suz12*/*Ezh2* (core components of the repressor complex *PRC2*), respectively. In addition, we demonstrate that compound pharmacological inhibition of Il3, Gm-Csf and Ezh2 restores normal hematopoiesis and colon homeostasis in a well-established preclinical model of colitis, establishing E2f as a critical driver of stress myelopoiesis.

## RESULTS

### Disrupting the Rb/E2f interaction alters the contribution of MPP4 cells to hematopoiesis

To determine the consequences of HSPC exit from quiescence, we previously used a mouse model that enables the conditional inactivation of *Rb* family members (*Rb^Lox/Lox^*, *p130^Lox/Lox^*, *p107^-/-,^* aka Triple Knock Out-*TKO*). Crossing *TKO* mice with *Mx1-Cre* and *Rosa26-CreER^T2^*transgenes, we first showed that derepression of E2f activity in hematopoietic cells is sufficient to recapitulate the features of stress myelopoiesis, including sustained HSPC proliferation and enforced myeloid-biased differentiation at the expense of lymphopoiesis^27,28^. A major advantage of the *TKO* model is the absence of background molecular noise that is inevitably generated upon stimulation of HSPCs by pleiotropic inflammatory mediators^29–32^, thereby enabling the identification of a mechanism strictly activated by HSPC exit from quiescence. Here, for experimental consistency, we have used the same mouse models to investigate the nature of this mechanism.

We first assessed the frequency of HSPC (aka KLS cells, which includes LT-HSC and MPP1-4) subpopulations in *CT* (*Rb^lox/lox^, p130^lox/lox^ p107^-/-^* used as controls) and *TKO* (*Rosa26-CreER^T2^ Rb^Lox/Lox^*, *p130^Lox/Lox^*, *p107^-/-^)* bone marrow (BM), two weeks after *Rb* family inactivation. Within the subpopulations, lymphoid-primed MPP4 frequency significantly increases while MPP1 frequency (aka short-term HSC, ST-HSC) decreases in *TKO* BM (**Fig. 1a** and **Fig. S1a**). Functionally, *ex vivo* culture assays show increased myeloid colony forming activity (**Fig. 1b**) and decreased lymphopoiesis (**Fig. 1c**) for *TKO* MPP1, MPP3 and MPP4 compared to their respective controls, suggesting a myeloid bias in multiple *TKO* HSPC subpopulations. Moreover, the fraction of MacI^+^ myeloid cells that have differentiated from *TKO* MPP3 and MPP4 when placed in liquid culture is greater than that of their *CT* counterparts, suggesting that the differentiation of MPP3 and MPP4 into mature populations is accelerated upon *Rb* family deficiency (**Fig. 1d** and **Fig. S1b**). To assess the cell intrinsic nature of this myeloid priming, we transplanted *Rag1^-/-^*immunodeficient mice with unfractioned *CT* or *cTKO* BM cells and treated the recipient mice with Tamoxifen five weeks after transplantation (when donor hematopoiesis is established). Immunophenotypic analysis of chimeric BM shows a progressive increase in *TKO* MPP4 frequency compared to *CT* MPP4 over time (**Fig. S1c**). In addition, colony forming activity assay shows increased production of granulocyte/monocyte (G/M/GM) colonies from *TKO* MPP4 compared to controls, showing that their expansion and accelerated myeloid differentiation is cell autonomous (**Fig. S1d**).

**Figure 1.**
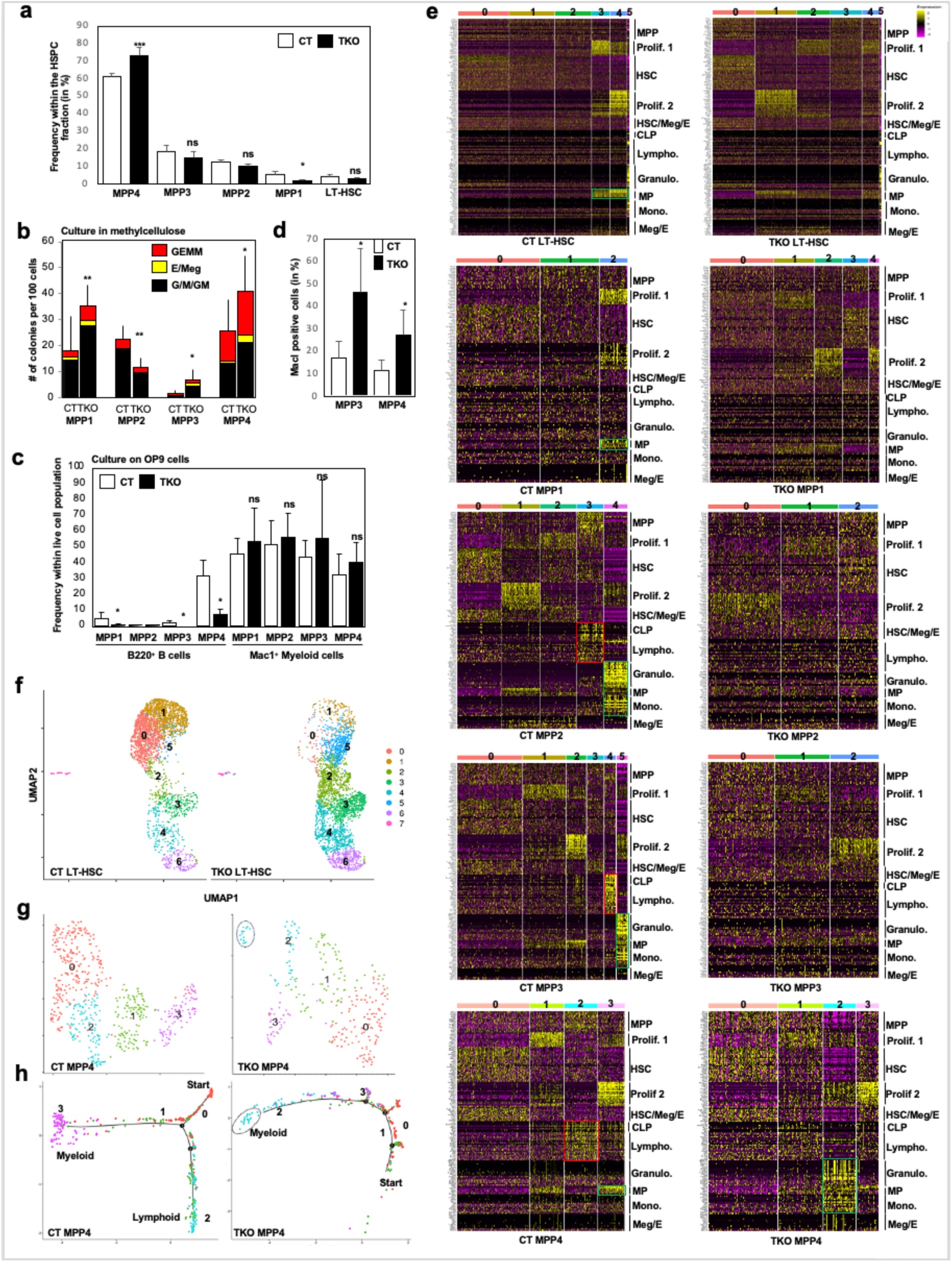
**a** Using surface markers that enable the identification of LT-HSC and MPP1-4 populations (Kit, Lineage Markers, Sca1, Cd48, Cd150, Flk2), flow cytometry analysis shows the frequencies for *CT* and *TKO* LT-HSC/MPP1-MPP4. *CT*, white bars; *TKO*, black bars. n=7. **b** 100 cells from each subpopulation were plated into standard methylcellulose. The number and identity of colonies were quantified after 8 days in culture (n=3). **c** 100 *CT* and *TKO* MPP1-MPP4 cells were co-cultured with OP9 cells and the number of B cells (Cd45^+^, B220^+^) and Myeloid cells (Cd45^+^, Mac1^+^) were quantified by flow cytometry after 10 days in culture (n=4). **d** *CT*/*TKO* MPP3 and MPP4 were isolated by flow cytometry and placed in liquid culture. After 5 days, surface expression of the pan-myeloid marker MacI was assessed in cultured cells to quantify their differentiation rate. **e** heatmap showing the expression of representative genes for different steps of hematopoiesis in *CT* and *TKO* LT-HSC, MPP1-4 subpopulations. **f** Individual sc-Seq datasets for *CT* and *TKO* LT-HSCs were merged for analysis and individually displayed to determine the distribution of clusters from the merged dataset per genotype. **g** UMAP display of sc-Seq analysis for *CT* (left) and *TKO* (right) MPP4. Cells surrounded by a dotted line represent a projected subcluster. **h** Trajectory analysis by Monocles-2D identifies a bipotential (myeloid/lymphoid) fate in *CT* MPP4 (left panel) while the lymphoid fate is obliterated in *TKO* MPP4 (right panel). Cells surrounded by a dotted circle represent a projected subcluster (i.e located at the end of the trajectory).

To characterize the consequences of *Rb* family inactivation on HSPC contribution to hematopoiesis at the single cell level, we performed 10x single-cell RNA-Sequencing (scRNA-Seq) on HSPC subpopulations (LT-HSC and MPP1-4) from *CT* and *TKO* mice. Based on the expression of an array of markers corresponding to the different steps of hematopoiesis (as determined by Gene Expression Commons^33^-see Material&Methods for gene list), we found that a myeloid signature (as determined by expression of markers for granulocytes, monocytes or myeloid progenitors-green box) is expressed in all *CT* subpopulations (**Fig. 1e**). On the other hand, the myeloid signature disappears in all *TKO* subpopulations except for *TKO* MPP4. While clusters 0, 1 and 3 in *CT* MPP4 display similar marker expression profile as clusters 0, 1 and 3 in *TKO* MPP4, cluster 2 in *CT* MPP4 displays a lymphoid signature (as determined by expression of markers for common lymphoid progenitor-CLP- and lymphocytes-red box) whereas cluster 2 in *TKO* MPP4 displays a myeloid signature that includes mature granulocytes and monocytes markers. In addition, the lymphoid signature is also present in *CT* MPP2 and MPP3 but disappears in all *TKO* MPP subpopulations.

Merging both *CT* and *TKO* LT-HSCs datasets into a single dataset reveals that the most primitive (based on HSC marker expression-**Fig. S1e**) clusters (0, 1, 2, 5) are asymmetrically enriched in *CT* (clusters 0&1) and *TKO* (cluster 2&5) LT-HSCs (**Fig. 1f**): cluster 2&5, which express a mix HSC/MPP marker signature (**Fig. S1e**), are located on the outer edge of the main trajectory (**Fig. S1f**), suggesting that they represent alternate primitive states that do not exist in homeostatic conditions. In addition, downstream (based on trajectory) clusters 3&4&6 are present in both *CT* and *TKO* LT-HSCs. However, the ratio of downstream (cluster3&4&6) vs primitive (cluster0&1&2&5) cells is higher in *TKO* versus *CT* LT-HSCs, suggesting that *Rb* family inactivation promotes the differentiation of the most primitive LT-HSCs.

Downstream of LT-HSCs, UMAP display and trajectory analysis shows that, in *CT* MPP4 cells, the more primitive cluster 0 differentiates into both lymphoid-primed cluster 2 and myeloid-primed clusters 1&3. In contrast, cluster 0 exclusively differentiates towards myeloid-primed cluster 2 cells in *TKO* MPP4 cells (**Fig. 1g-h**) while the lymphoid fate is absent. Collectively, these data suggest that *Rb* family inactivation expands MPP4 population and primes its differentiation exclusively towards myelopoiesis to accelerate the production of myeloid cells.

### Accelerated differentiation of *TKO* HSPCs towards the granulocyte/monocyte lineage

MPPs differentiate into lineage-committed myeloid progenitors (MPs, found in the Lin^-^ Kit^+^ Sca1^-^ fraction), which consist of distinct progenitor subpopulations with progressive lineage commitments to either megakaryocyte/erythrocyte (MEP) or granulocyte/monocyte (GMP) lineages. We previously shown that *Rb* family inactivation increases GMP frequency at the expense of CMP and MEP frequency^28^ and liquid culture confirms that *TKO* MP cells differentiates faster than their *CT* counterparts (**Fig. S2a**). Using a set of markers that enables a more thorough identification of MP subpopulations along the progressive granulocyte and monocyte commitment^34^, we found that *Rb* family inactivation increases the frequency of CMP- Flt3^-^ cells while decreasing the frequency of MDPs (monocyte/dendritic progenitors), suggesting a biased differentiation of the upstream *TKO* CMP-Flt3^+^ Cd115^lo^ subpopulation towards the granulocytic fate (**Fig. 2a**). However, *ex vivo* colony forming assay does not identify increased cellular output for *TKO* CMP-Flt3^+^ Cd115^lo^ and CMP-Flt3^-^ cells compared to their respective controls (**Fig. 2b**), suggesting that the enhanced production of myeloid cells observed in *TKO* hematopoiesis is not driven by *TKO* MP cells.

**Figure 2.**
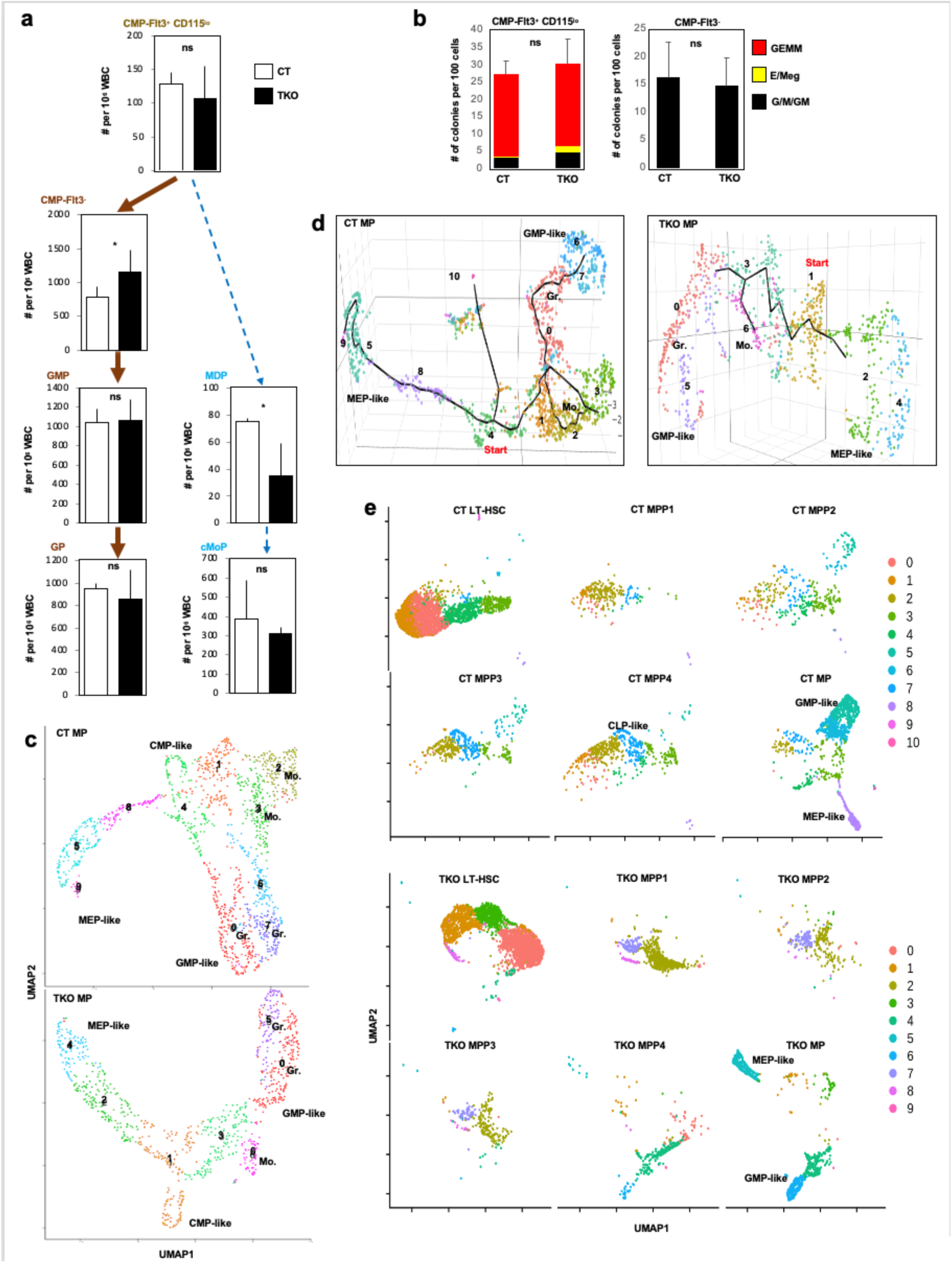
**a** Immunophenotypic analysis of MP subpopulations in *CT* and *TKO* BM two weeks after Tamoxifen treatment using antibodies against Kit, Fcγr, Cd34, Ly6c, Flt3 and Cd115 (n=3). Bold brown arrow indicates a differentiation pathway that is dominant in *TKO* mice compared to the alternative pathway that is marked by a blue dotted line. **b** Colony assay for 100 *CT*/*TKO* CMP-Flt3^+^ Cd115^lo^ (left) and CMP-Flt3^-^ (right) cells plated in standard methylcellulose (n=3). **c** UMAP display of *CT* (1,979 cells) and *TKO* (1,149 cells) MPs. CMP-like, GMP-like and MEP- like are identified based on gene expression in the heatmap. **d** Trajectory analysis by Monocles- 3D of *CT* and *TKO* MP cells. Start is identified as the cluster expressing the highest level of a LT-HSC signature. **e** sc-Seq datasets for *CT* (upper panels) and *TKO* (lower panels) LT-HSC, MPP1-4 and MP were merged for analysis and individually displayed by UMAP to determine the distribution of each population within the hematopoietic progenitor compartment.

To characterize the composition of the MP compartment and investigate its integration with upstream MPP progenitors upon *Rb* family inactivation, we performed scRNA-Seq on *CT* and *TKO* MP cells isolated from the BM two weeks after Tamoxifen treatment. The number of clusters with a myeloid-like signature is decreased from five clusters in *CT* MPs (clusters 0, 2, 3, 6 and 7) to three clusters (clusters 0, 5 and 6) in *TKO* MPs (**Fig. 2c**), with a particular reduction of the number of cells with a monocytic signature (**Fig. S2b**). Trajectory analysis shows that the differentiation of *CT* MPs includes a short linear path from the most primitive cluster (cluster 4) to MEP-like cells (clusters 5, 8 and 9, based on expression of an erythrocyte signature-**Fig. S2b**) and a long and circumvolved path to GMP-like cells (**Fig. 2d**). In contrast, *TKO* MP differentiation is characterized by a short linear path from the most primitive cluster (cluster 1) that rapidly diverges into either a granulocyte path (clusters 0&5) or a monocyte path (cluster 6). These data further suggest that *Rb* family inactivation alters the differentiation paths of early MP subpopulations to accelerate myelopoiesis.

Finally, we merged individual LT-HSC, MPPs and MP datasets into a single dataset per genotype. In both *CT* and *TKO* conditions, an upstream immature population (clusters 0, 1 and 2 for *CT*; clusters 0, 2 and 3 for *TKO*) diverges into either MEP or GMP fates (**Fig. S2c-d**); however, this immature population displays increased *Ki67* expression upon *Rb* family inactivation compared to CT (**Fig. S2e**). UMAP visualization of individual HSPC subpopulations within these merged datasets shows that, while MPP4 cells overlap with other MPP cells in *CT* mice, *TKO* MPP4 cells predominantly overlap with *TKO* MP cells engaged in the GMP-like pathway (**Fig. 2e**), confirming their enhanced commitment to myelopoiesis.

Together, these data identify the expansion of MPP4 and their accelerated differentiation towards myelopoiesis as a dominant feature of hematopoietic progenitors upon *Rb* family inactivation.

### A limited number of transcriptional networks drive *TKO* myelopoiesis

Analysis of transcriptional network activity with SCENIC^35^ identifies 190 active transcription factors in lymphoid-primed *CT* MPP4 cluster 2, suggesting their uncommitted fate. In contrast, only 40 transcription factors are active in myeloid-primed *TKO* MPP4 cluster 2, including *E2f* and a network of monocytic/granulocytic transcription factors (*Irf8*, *Cebp* family, *Xbp1*, *Spi1*) (**Fig. 3a**).

**Figure 3.**
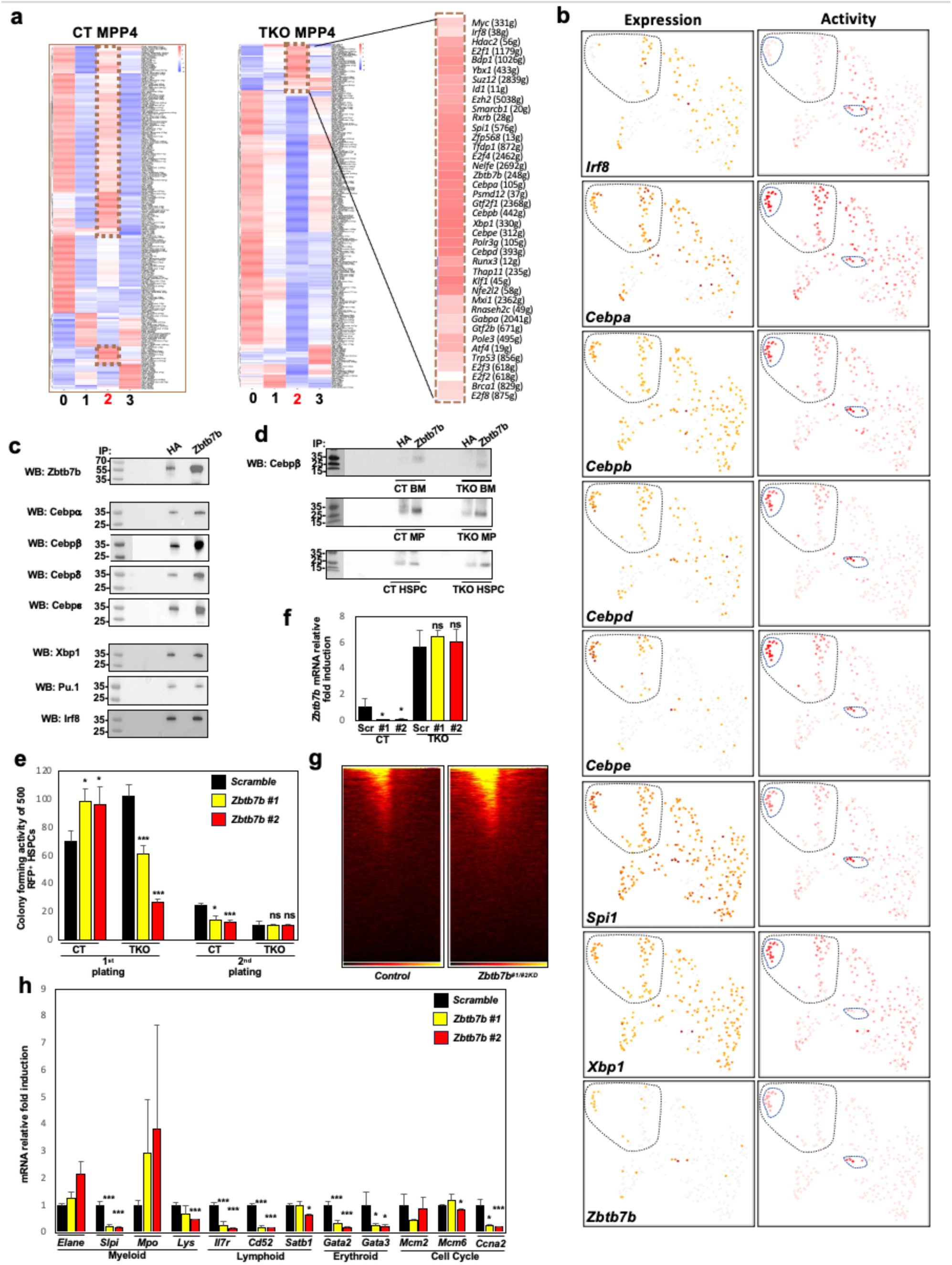
**a** Heatmap display of SCENIC analysis identifies transcriptional networks active in each cluster from *CT* and *TKO* MPP4. Networks active in *CT* cluster 2 (lymphoid-primed, 190 networks) and *TKO* cluster 2 (myeloid-primed, 40 networks) are highlighted with a dashed brown rectangle. Transcriptional networks active in *TKO* MPP4 cluster 2 are magnified at the right, with the network in *italic* and the number of target genes (g) activated by this network in cluster 2. **b** Expression (left) and activity (right) level assessed by SCENIC for a myeloid network (monocytic: *Irf8*. granulocytic: *Cebpa/b/d/e*, *Spi1* and *Xbp1*) as well as *Zbtb7b* in TKO MPP4 are displayed at the single cell level to highlight their overlapping pattern. Cell clusters with active *Zbtb7b* are circled with a blue dotted line and these circles are applied to the myeloid network factors for better comparison. Cells corresponding to cluster 2 (myeloid-primed) are circled with a black dotted line. Color code: Pale: low expression/activity. Dark(er): high(er) expression/activity. **c** Protein lysis from *CT* BM cells were equally split and immunoprecipitated with polyclonal antibodies against either HA (negative control) or Zbtb7b. The presence of myeloid transcription factors in the precipitate was assessed by Western Blotting. Each factor was tested in independent experiments to avoid potential membrane reblotting issues. **d** Identical number of *CT* and *TKO* cells from unfractioned Bone Marrow (BM), flow-isolated Myeloid Progenitor (MP, Lineage^-^, Kit^+^, Sca1^-^) and HSPC fractions were isolated two weeks after Tamoxifen-induced *Rb* family loss. Protein lysis extracts were equally split and immunoprecipitated with polyclonal antibodies against either HA (negative control) and Zbtb7b. The presence of Cebpβ in the precipitated fractions was assessed by Western Blotting. **e** Colony forming activity of 500 RFP^+^ *CT* and *TKO* HSPC cells infected with lentiviruses expressing either a scramble hairpin or two hairpins (#1/#2) against *Zbtb7b*. Colonies were counted after 9 days in culture (1^st^ plating). On the same day, 10,000 colony cells were replated and their colony forming activity was reassessed after 9 days (2^nd^ plating) (n=3). **f** *Zbtb7b* expression level in *CT* and *TKO* colony cells from the 2^nd^ plating, as determined by RT-qPCR (n=3). **g** TSS map from ATAC-Seq performed on *CT* and *TKO* colony cells from the 1^st^ plating shows opening of chromatin upon *Zbtb7b* silencing. **h** Expression of markers for different hematopoietic lineages was assessed by RT-qPCR in *CT* colony cells collected after the 2^nd^ plating (n=3).

Visualization of SCENIC analysis at the single cell level (**Fig. 3b**) reveals that the expression of the monocytic factor *Irf8* is mosaic in *TKO* MPP4 while its activity decreases along *TKO* MPP4 differentiation trajectory (as identified in **Fig. 1h)**. In parallel, the expression of granulocytic factors *Cebpa/b/d/e*, *Spi1* (aka *Pu.1*) and *Xbp1* progressively increases along the same trajectory, with their activity reaching its maximum in the projected subcluster of cluster 2 (see **Fig. 1h**). Among the 40 transcription factors active in *TKO* MPP4 cluster 2, *Zbtb7b* displays a pattern of transcriptional activity that uniquely overlaps with the pattern of granulocytic network activity (**Fig. 3b**, lowest panel). In contrast, *Zbtb7b* is not expressed in *CT* MPP4 (**Fig. S3a**), suggesting an earlier activation of its expression in *TKO* hematopoiesis. *Zbtb7b* (aka *Zfp67* or *ThPok*) is a zinc finger repressor protein that acts as a master regulator of lineage commitment via its chromatin remodeling activity. While mostly studied for its role in *Cd4^+^* T cell specification^36^, *Zbtb7b* is also expressed in the myeloid lineage^37^ and in particular granulocytes (**Fig. S3b**), suggesting a functional role in myelopoiesis. Immunoprecipitation of unfractioned *CT* BM cells shows that, among granulocytic factors, *Zbtb7b* only directly binds to Cebpβ, Cebpδ and Cebpε (**Fig. 3c**). Although the interaction between Zbtb7b and Cebpβ is observed in both *CT* and *TKO* MPs, it can only be observed in *TKO* HSPCs but not in their control counterparts, further suggesting that *Zbtb7b* expression occurs earlier in *TKO* hematopoiesis compared to normal hematopoiesis (**Fig. 3d**). Functionally, *Zbtb7b* silencing impairs the colony forming activity of *TKO* HSPCs, but not *CT* HSPCs, upon their first plating (**Fig. 3e-f**), opens up chromatin (**Fig. 3g**) and decreases the expression of mature markers for the principal hematopoietic lineages (**Fig. 3h**). Collectively, these results suggest that *TKO* myelopoiesis is driven by a limited transcriptional network that includes granulocytic transcription factors and *Zbtb7b*.

### E2f activity enhances βcytokine signaling via activation of their common βchain receptor

To determine whether granulocytic transcription factors and *Zbtb7b* share a common mechanism of transactivation in *TKO* HSPCs, we mined Encode database and found that *Stat3/5* are the only transcription factors with binding sites in all regulatory regions of *Zbtb7b*, *Cebp* family, *Spi1* and *Xbp1* (**not shown**). Accordingly, *Stat* signaling activity is enhanced in *TKO* MPP4 compared to *CT* (**Fig. S4a**). To identify the receptor signaling driving enhanced *Stat* activity in *TKO* HSPCs in an unbiased manner, we screened for the expression of 256 markers at the surface of *CT* and *TKO* hematopoietic progenitors (HSPCs and MPs). Among *Stat* signaling associated receptors, only Cd131 (the common βchain receptor for Il3rα and GM-Csfrα) is overexpressed in *TKO* HSPCs and MPs compared to their respective controls (**Fig. 4a; Fig. S4b-c**). In contrast, receptors for inflammatory cytokines such as *Il1r*, *Il6r*, *Tnfr1/2* and *Ifngr* are all decreased in *TKO* HSPCs compared to controls. Analysis of our array data^27,28^ confirms increased abundance of *Csf2rb* (which codes for Cd131) transcripts in *TKO* LT-HSCs and the more inclusive *TKO* HSPCs, compared to their respective controls (**Fig. S4d**). Further analysis at the single cell level confirms higher *Csf2rb* expression in *TKO* vs *CT* hematopoietic progenitors, in particular the myeloid-primed cluster 2 of *TKO* MPP4s (**Fig. S4e-h**). Sequence analysis identifies an evolutionarily conserved E2f binding site in mouse and human *Csf2rb* promoter regions and ChIP assays performed in unfractioned primary BM white blood cells confirm that E2f1 and E2f3 bind to *Csf2rb* promoter region (**Fig. 4b**), establishing *Csf2rb* as a novel non-canonical E2f target gene in hematopoiesis. Phospho-flow (p-flow) assay shows increased phosphorylation of Erk (a read-out for βcytokine signaling activity) in *TKO* HSPCs in response to stimulation by both Il3 and Gm-Csf compared to *CT*, confirming that increased surface expression of Cd131 is associated with its enhanced intracellular signaling (**Fig. 4c**). In support of our initial hypothesis, we found that stimulation of *CT* MPP4 with Il3 transactivates *Cebpb* and *Zbtb7b* expression, confirming that the granulocyte network and *Zbtb7b* are indeed regulated by βcytokine signaling (**Fig. 4d**).

**Figure 4.**
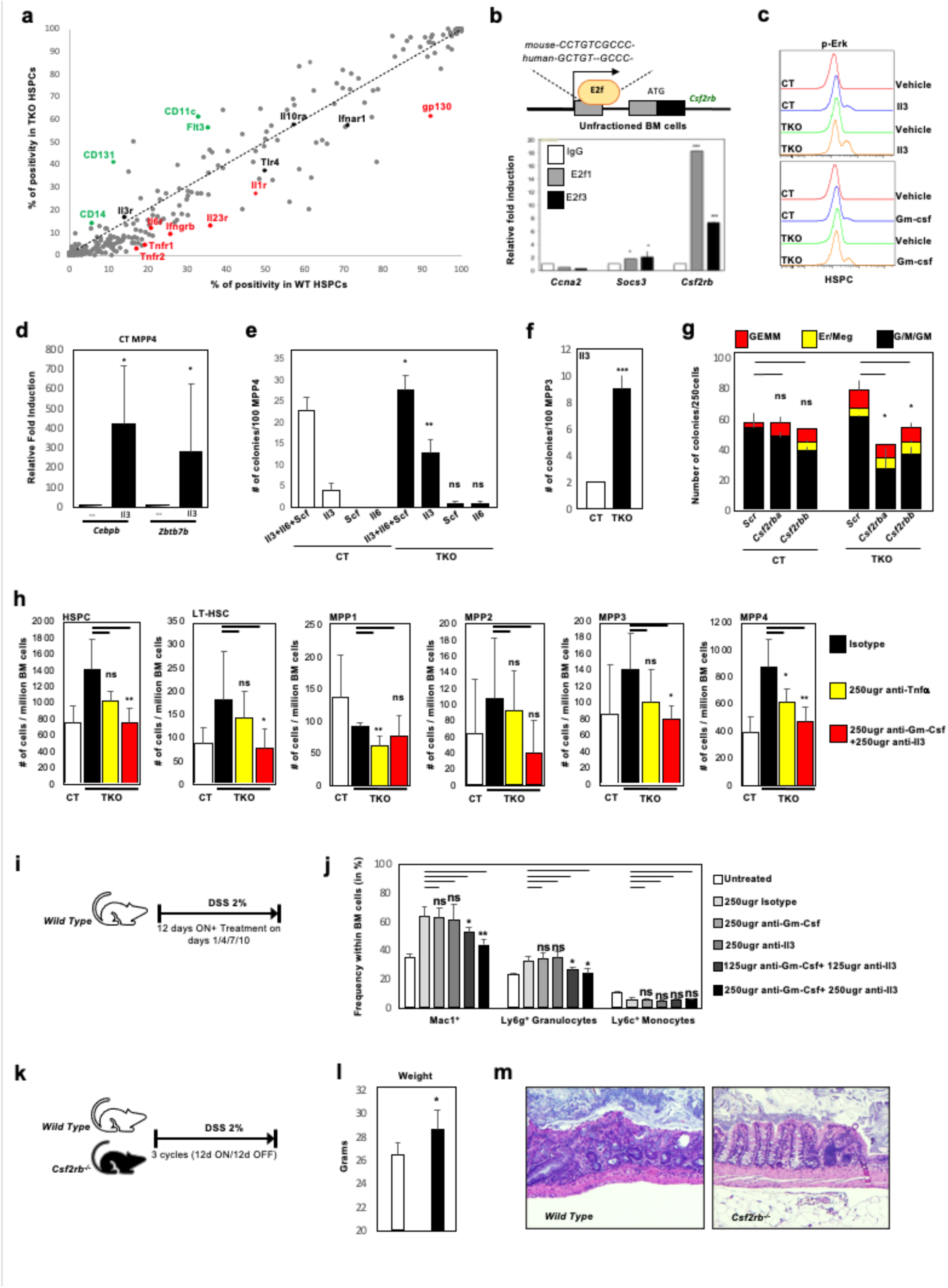
**a** Unfractioned BM cells from *CT*/*TKO* mice (5 per group) were isolated two weeks after Tamoxifen treatment, dyed for multiplexing purposes, merged, stained with antibodies against lineage markers (B220, Mac1, Gr1, Ter119, Cd3, Cd4, Cd8), Kit and Sca1 (HSPC) and screened for the expression of 256 surface markers (Biolegend antibody array® complemented by individual analysis for CD131 since it is not included in the array design). Background signal intensity was assessed by isotype staining. Each dot represents a surface marker expression in *CT* (x-axis) and *TKO* (y-axis) HSPCs. The dotted line represents identical positivity % in *CT* and *TKO* HSPCs. Markers of monocyte lineage (CD14 and CD11c) and MPP4 (Flt3) are highlighted for technical validation. Gene names of interest are color-coded for better visualization when expression difference between *CT* and *TKO* HSPCs is at least 10 points. **b** Upper panel: analysis of mouse and human *Csf2rb* regulatory region identifies a conserved E2f binding site in the first untranslated exon. Lower panel: ChIP analysis performed on unfractioned BM cells show that E2f1 and E2f3 bind to a region of the *Csf2rb* promoter that harbors the putative E2f binding site in primary BM cells (n=3). **c** *CT* and *TKO* HSPCs were isolated two weeks after Tamoxifen treatment, serum starved and stimulated with Il3 or Gm-Csf (10ng/ml) for 10 min. p-Erk was used as a readout for βcytokine signaling activity (n=3). **d** *CT* MPP4 were isolated by flow cytometry, serum starved for 2 hours and treated with 20ng/ml of Il3 for 1 hour. Cells were collected and RNA was extracted prior to qPCR to detect *Cebpb* and *Zbtb7b* expression. **e** 100 *CT* and *TKO* MPP4 cells were plated in cytokine-free methylcellulose, supplemented with either a cocktail of Il3, Il6 and Scf, or Il3, Scf and Il6 only. Colonies were counted after 9 days in culture (n=3). **f** 100 *CT* and *TKO* MPP3 cells were plated in cytokine-free media, supplemented with Il3. Colonies were counted after 9 days in culture (n=3). **g** *CT* and *TKO* MPP4 cells were infected with p-Sicor- RFP lentiviral vectors expressing either a scramble hairpin or two hairpins (*a* and *b*) targeted against *Csf2rb*. 500 RFP^+^ cells were plated for methylcellulose culture and colonies were quantified after 9 days in culture (n=3). **h** *CT* and *TKO* mice were treated with Tamoxifen for four consecutive days. On day five, eight and eleven, *TKO* mice were injected with either isotype antibody (control, black bar), antibody against Tnfα (250ug, yellow bar) or a combination of antibodies against Gm-Csf and Il3 (250ug each, red bar). On day 14, HSPC subpopulations were analyzed for frequency per million BM cells (n=6 per groups). **i** DSS 2% was added to the drinking water of wild-type mice for 12 days. Mice were injected with various combinations of blocking antibodies (isotype, anti-GM-Csf, anti-Il3, combination of anti-GM-Csf and anti-Il3) on day 1, 4, 7 and 10 of colitis-inducing DSS treatment. Untreated wild-type mice were used as control. **j** Frequency of Mac1^+^ cells, Ly6g^+^ granulocytes and Ly6c^+^ monocytes in the BM. **k** Wild-type and *Csf2rb* deficient mice were exposed to three cycles of DSS 2% treatment (1 cycle: 12 days with DSS 2% containing drinking water followed by 12 days with normal drinking water). **l** Mice were weighted at the end of the treatment. **m** Representative H&E staining of intestine sections from *wild-type* and *Csf2rb-/-* mice shows more severe alteration of villae structure in wild-type mice compared to *Csf2rb* deficient mice (n=5).

Functionally, we found that *TKO* MPP4 as well as *TKO* MPP3 are hyperresponsive to Il3 stimulation compared to their respective *CT* counterparts *ex vivo* (**Fig. 4e-f; Fig. S7i**). On the other hand, *TKO* LT-HSCs are not hyperresponsive to Il3 stimulation while *TKO* MPs are hyperresponsive to βcytokines (**Fig. S7j-k**), suggesting that most *TKO* progenitor subpopulations display increased βcytokine signaling except for primitive *LT-HSCs*. *Ex vivo*, knockdown of *Csf2rb* expression is sufficient to repress the colony forming activity of *TKO* MPP4 down to *CT* levels (**Fig. 4g**). *In vivo*, treatment of *TKO* mice with blocking antibodies for Gm-Csf and Il3 is more potent than TNFα blockade to restore HSPC frequency to *CT* level (**Fig. 4h**). Collectively, these data suggest that E2f enhances βcytokine signaling via direct transactivation of *Csf2rb* to activate a transcriptional network that promotes *TKO* myelopoiesis. Review of literature shows that increased Cd131 expression at the surface of HSPCs is observed in various inflammatory contexts, suggesting that enhanced βcytokine signaling is a conserved feature of stress myelopoiesis^38–46^.

Although *Csf2rb* deficiency does not alter BM cellular composition in homeostatic conditions (**Fig. S5a-b**), it decreases the *ex vivo* colony forming activity of MPP4 cells compared to their control counterparts (**Fig. S5c**), further suggesting a critical role for βcytokine signaling in the MPP4 subpopulation. To determine the consequences of targeting βcytokine signaling in relevant inflammatory contexts, we subjected wild-type and *Csf2rb* deficient mice to two different types of stress. In a first inflammatory model, addition of Dextran Sodium Sulfate (DSS) to the drinking water initiates a well-documented model of colitis^47^. Exposure of wild type mice to 2% DSS for 12 days increases myeloid cell frequency in the BM, which is decreased upon treatment with a combination of individual blocking antibodies against Il3 and Gm-Csf (**Fig. 4i-j**). Moreover, *Csf2rb* deficiency prevents weight loss and improves intestinal architecture in the context of chronic exposure to DSS (**Fig. 4k-m**). In a second inflammatory model, genetic and pharmacological inhibition of βcytokine signaling represses the accumulation of granulocytes and improved intestinal and skin architecture upon exposure to LPS (**Fig. S5d-i**), a pro-inflammatory molecule that mimics bacterial infection. Overall, these data suggest that targeting βcytokine signaling activity decreases stress myelopoiesis and improves clinical conditions in inflammatory contexts.

### Activation of *Ezh2*/*Suz12* by E2f represses alternative cell fates during *TKO* myelopoiesis

Decreased production of alternative hematopoietic lineages such as erythrocytes and lymphocytes is a hallmark of stress myelopoiesis. Reintroduction of a single *p107* allele in the *TKO* background represses excessive myelopoiesis without restoring normal lymphopoiesis^28^, suggesting that the repression of alternative lineages may be driven by a distinct mechanism during stress myelopoiesis. ATAC-Seq analysis shows that the chromatin structure of *CT* and *TKO* MPP4s is overall similar, with a limited number of genomic regions uniquely open either in *CT* MPP4 (749 loci) or in *TKO* MPP4 (94 loci) (**Fig. 5a-b; Fig. S6a**). Motif analysis of the 749 genomic regions open in *CT* MPP4 and closed in *TKO* MPP4 reveals an enrichment for transcription factors that have documented roles in stemness maintenance as well as erythropoiesis (**Fig. 5c**). SCENIC analysis shows that transcriptional networks active in promoting cell fate alternatives to myelopoiesis such as erythropoiesis, lymphopoiesis and stemness are progressively repressed along *TKO* MPP4 trajectory (**Fig. 5d**; **Fig. S6b**), overall suggesting that alternative hematopoietic lineages are actively repressed during stress myelopoiesis.

**Figure 5.**
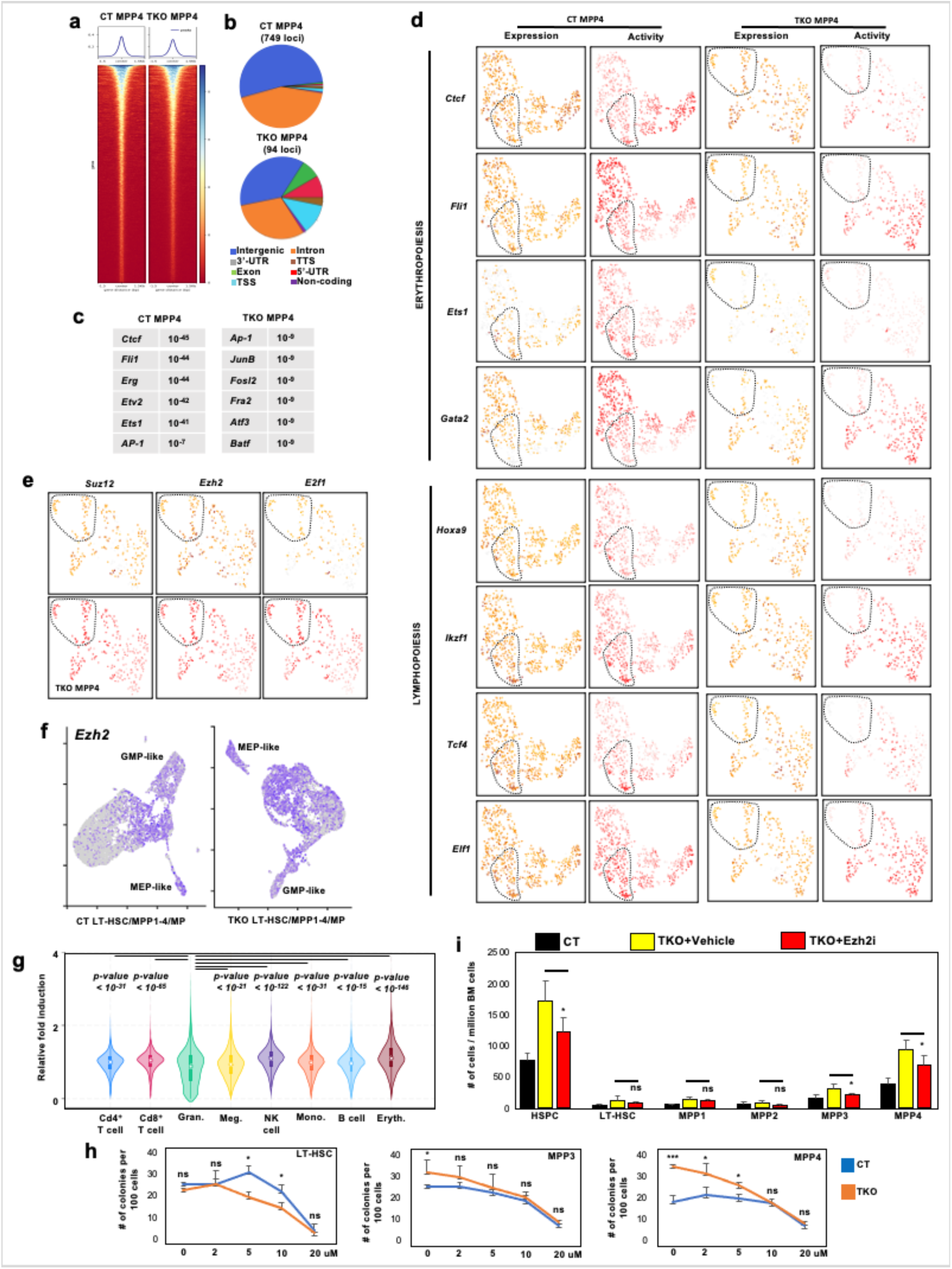
**a** heatmap for the ATAC-Seq analysis of *CT* and *TKO* MPP4 cells isolated two weeks after *Rb* family inactivation. The upper panels indicate the number (y axis) of opened region based on their length (x axis) for each population. **b** pie chart display of opened chromatin regions (peaks) that are unique to either *CT* or *TKO* MPP4, based on their genomic location. **c** Top results for motifs enrichment analysis for chromatin regions that are unique to either *CT* or *TKO* MPP4 identifies motifs for erythropoiesis-associated transcription factors in *CT* MPP4. **d** SCENIC analysis of the expression and activity of representative transcriptional networks involved in the the promotion of erythropoiesis and lymphopoiesis in *CT* and *TKO* MPP4. **e** SCENIC analysis for the expression (up) and activity (down) for Suz12, Ezh2 and E2f1 in *TKO* MPP4. **f** *Ezh2* expression at the single cell level in the merged *CT* (left) and *TKO* (right) LT-HSC/MPP1-4/MP datasets as identified by Seurat. **g** Statistical analysis of 5038 *Ezh2* target gene (as identified by SCENIC in **Fig. 3a**) expression profile through the different mature hematopoietic populations, as determined by Gene Expression Commons. **h** Colony forming activity of 100 *CT* and *TKO* LT-HSC (left), MPP3 (middle) and MPP4 (right) cells in the presence of increasing amount of FDA-approved Tazemetostat®, a selective *Ezh2* inhibitor. **i** *CT* and *TKO* mice were treated with Tamoxifen for four days. *TKO* mice were subsequently either treated with vehicle or received 5mg/day of FDA- approved Tazemetostat® by gavage. On day 14, HSPC subpopulations were analyzed by flow cytometry. Number of cells for each subpopulation per million BM cells (n=6 per groups).

Screening through transcriptional networks that are active in myeloid-primed cluster 2 of *TKO* MPP4 (**Fig. 3a**), we found a unique inverse correlation between the progressive repression of these alternative cell fate networks and the progressively increased expression and activity of Ezh2 and Suz12, two core members of the PRC2 repressive complex and E2f target genes^48,49^, along *TKO* MPP4 trajectory (**Fig. 5e**). Accordingly, *Ezh2* expression is increased in *TKO* progenitors versus their *CT* counterparts (**Fig. 5f**). SCENIC identifies 5038 *Ezh2* target genes in myeloid-primed cluster 2 of *TKO* MPP4 (**Fig. 3a**), suggesting a significant role for *Ezh2* in shaping *TKO* hematopoiesis. Expression analysis for these 5038 genes across individual mature hematopoietic lineages (using Gene Expression Commons as a data source) shows that these *Ezh2* target genes are significantly repressed in granulocytes compared to all other lineages, establishing a correlation between their reduced expression and commitment to the granulocytic fate (**Fig. 5g**; **Fig. S6c**).

To functionally determine the role of Ezh2 in *TKO* myelopoiesis, we favored a pharmacological approach to inhibit its activity, since its genetic ablation in the hematopoietic compartment initiates malignancies of various natures^50,51^ that would complicate the interpretation of our experiments. First, we found that the *ex vivo* colony forming activity of various *TKO* HSPC subpopulations is more sensitive to Ezh2 inhibition generated upon gradual addition of FDA-approved Tazemetostat® than their *CT* counterparts (**Fig. 5h**). *In vivo*, Tazemetostat® represses *TKO* HSPC frequency, in particular within the *TKO* MPP3 and MPP4 populations, identifying a role for *Ezh2* in the regulation of *TKO* hematopoiesis (**Fig. 5i**). Collectively, these data suggest that *E2f* activates core members of the PRC2 repressive complex in *TKO* HSPCs to repress alternative hematopoietic lineages.

Next, we sought to determine the consequences of simultaneously targeting βcytokine signaling and Ezh2 activity for *TKO* myelopoiesis. In an experimental setting similar to **Fig. 4h** and **Fig. 5i**, dual treatment represses *TKO* HSPC cell frequency down to *CT* level (**Fig. 6a-b**). Cell cycle analysis shows that dual treatment does not decrease the frequency of *TKO* MP and MPP4 cells in the S phase of cell cycle (**Fig. 6c**), suggesting that βcytokine signaling and Ezh2 only modulate the differentiation of TKO HSPCs without controlling their proliferation. Next, we performed additional *ex vivo* experiments to rule out a non-cell autonomous consequence of Ezh2 inhibition on *TKO* HSPC differentiation. In a colony forming assay stimulated by either Il3 or Gm-Csf, dual treatment generates an additive effect on the growth inhibition of *CT* BM cells (**Fig. 6d**). In another colony forming assay stimulated by a standard cytokine cocktail, dual Il3 (Gm-Csf is not present in the M3434® media) and Ezh2 inhibition also generate an additive inhibitory effect on the colony forming activity of unfractioned *TKO* BM cells, but not their *CT* counterparts, suggesting that *TKO* hematopoiesis is particularly sensitive to dual inhibition (**Fig. 6e**). Analysis of colony cell lineage marker expression by flow cytometry reveals that single Ezh2 inhibition derepresses the formation of erythroid cells while single Il3 inhibition modestly represses the formation of mature myeloid cells. However, dual treatment achieves myeloid cell repression and erythroid expansion (**Fig. S7**). Based on these data, we propose a model where E2f coordinates the activation of its canonical cell cycle program with the non-canonical activation of βcytokine signaling and Ezh2 to drive *TKO* myelopoiesis (**Fig. 6f**).

**Figure 6.**
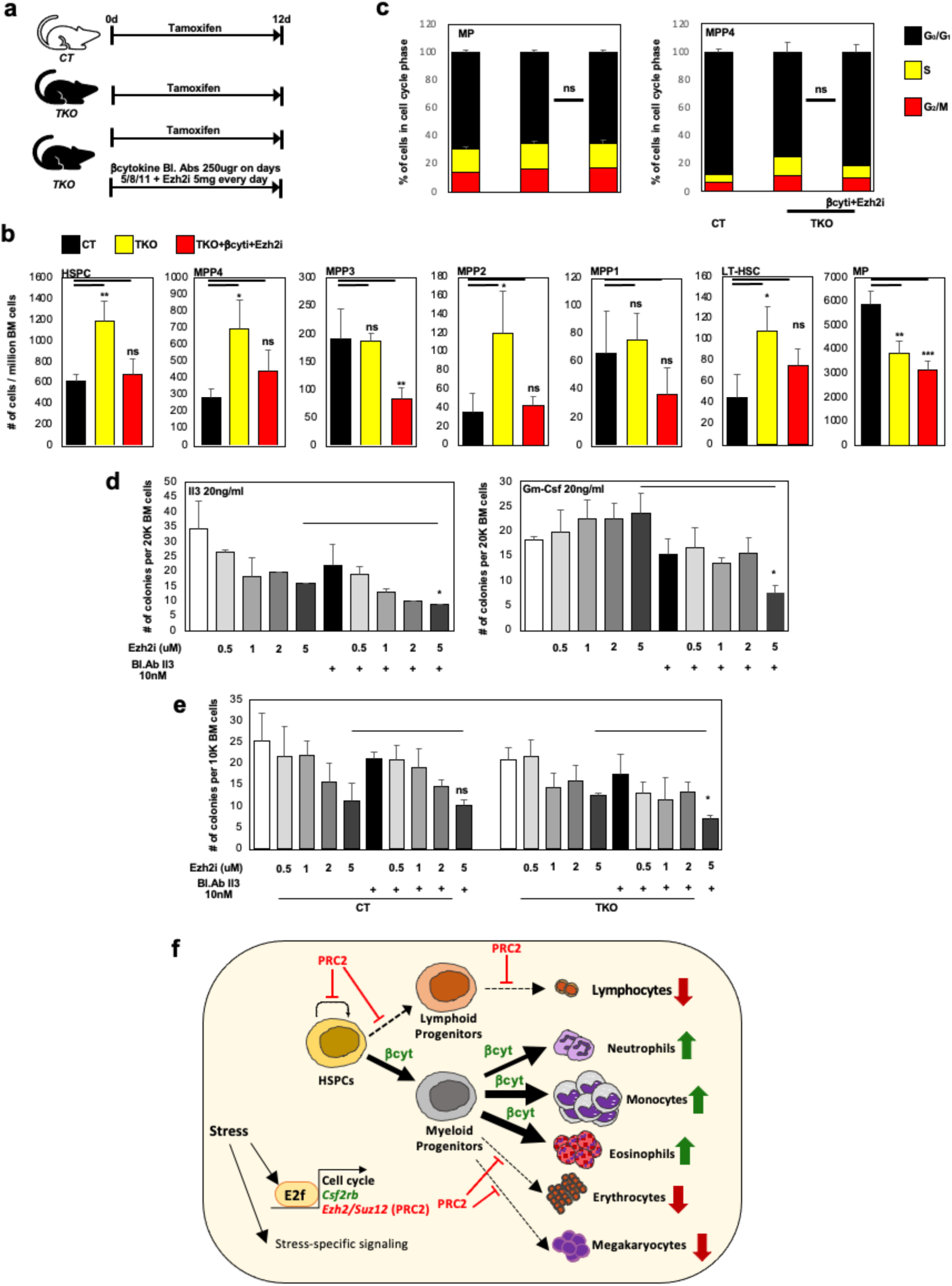
**a** *CT* and *TKO* mice were treated with Tamoxifen. Half of *TKO* mice were subsequently injected with 250ug of blocking antibodies against Gm-Csf and Il3 on days 5/8/11 while treated with 5mg of Tazemetostat® every day (n=4/group). **b** On day 12, HSPC and MP subpopulations were analyzed by flow cytometry. Number of cells for each subpopulation per million BM cells (n=4 per groups). **c** MP and MPP4 cells were isolated on day 12 from mice exposed to the different regimens and assessed for their cell cycle activity by PI incorporation (n=3). **d** 20K *CT* BM cells were plated in cytokine-free methylcellulose media supplemented with either Il3 (left; 20ng/ml) or Gm-Csf (right; 20ng/ml). Cells were treated with increasing dose of FDA-approved Tazemetostat® and/or blocking antibody against the respective βcytokine. Colonies were counted after nine days in culture (n=3). **e** 10K *CT* and *TKO* BM cells were plated in cytokine-containing methylcellulose (M3434®), together with increasing amount of FDA approved-Tazemetostat® and/or blocking antibody against Il3. Colonies were counted after nine days in culture (n=3). **f** Model: External stimuli of various natures such as inflammatory cytokines, growth factors, etc disrupt the inhibitory interaction between Rb and E2f. As a consequence to these stress signals, E2f combines the activation of its canonical cell cycle program with the non-canonical activation of *Csf2rb* and *Suz12/Ezh2* expression. While increased *Suz12*/*Ezh2* expression represses alternative hematopoietic fates (erythropoiesis, lymphopoiesis, stemness), increased *Csf2rb* promotes the production of inflammatory myeloid cells by enhancing Il3 and Gm-Csf signaling. In this model, the E2f-driven internal response is fine-tuned by parallel, stress-specific signaling activity.

### Dual βcytokine signaling and Ezh2 inhibition represses inflammatory conditions

To determine the consequences of dual inhibition for stress myelopoiesis in a clinically relevant pre-clinical inflammatory context, we further took advantage of the DSS-induced model of colitis introduced in **Fig. 4**. *CT* mice were exposed to DSS1% for 10 days and treated with various drug regimens to achieve either βcytokine, Ezh2 or dual βcytokine/Ezh2 inhibition (**Fig. 7a**). In the BM, DSS treatment triggers expansion of all hematopoietic progenitor subpopulations, except for MPP1 cells. Although single inhibition of either βcytokine signaling or Ezh2 only partially represses progenitor expansion, dual inhibition restores progenitor cell frequency to normal level (**Fig. 7b-c**). Functionally, dual inhibition represses the colony forming activity of BM cells as well as splenocytes back to normal level (**Fig. 7d**), without altering the cell cycle activity of MP and HSPC cells isolated from the BM (**Fig. 7e**). Within the mature cell compartment, Mac1^+^ myeloid cell frequency (**Fig. 7f**) and expression of key inflammatory markers (**Fig. 7g**) is also restored to normal level upon dual inhibition. Outside of the BM, dual inhibition similarly decreases the presence of myeloid cells in the colon (**Fig. S8a**). However, the inflammatory status of both granulocytes and monocytes (based on Cd18 and CCR2 surface expression, respectively) present in the colon as well as different biological compartments remains unaltered upon dual inhibition (**Fig. 7h-i**). Collectively, these data suggest that dual inhibition represses colitis-triggered stress myelopoiesis. At the initial site of inflammation, histological analysis of the colon by a veterinary pathologist confirms that exposure to DSS leads to increased infiltration of inflammatory cells as well mucosal ulceration. While single βcytokine signaling or Ezh2 inhibition only leads to modest improvement, dual inhibition decreases inflammatory cell infiltration and restores normal colon architecture in 80% of treated mice (**Fig. 7j**).

**Figure 7.**
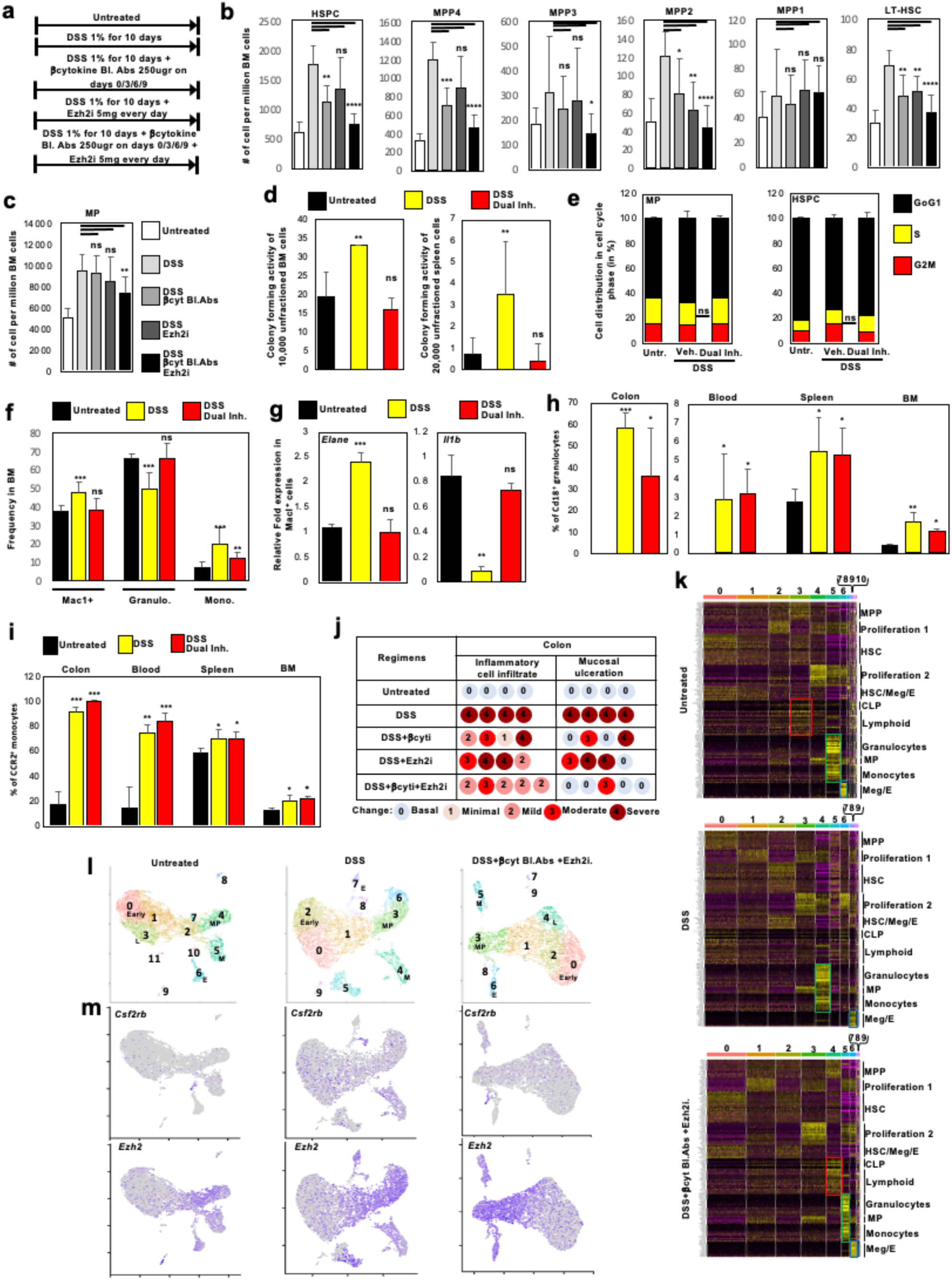
**a** *CT* mice were either untreated or treated with DSS1% in the drinking water for 10 days, with or without treatment to block βcytokine signaling (250ug of Il3 blocking antibody and 250ug of Gm-Csf blocking antibody on days 0, 3, 6 and 9 of DSS treatment) and/or treatment to block Ezh2 activity (Tazemetostat®, 5mg/day). **b-c** Analysis of HSPC subpopulation (**b**) and MP cell (**c**) frequency in the BM of *CT* mice treated with different regimens (n=6 per group). **d** Colony forming activity of 10,000 unfractioned BM cells (left) and 20,000 unfractioned spleen cells (right) isolated from a pool of mice treated with different regimens (n=3). **e** MP and HSPC cells were isolated from mice exposed to the different regimens and assessed for their cell cycle activity by PI incorporation (n=4). Untr: Untreated. **f** Analysis of mature myeloid populations (Mac1^+^), Granulocytes (Ly6g^+^) and Monocytes (Ly6c^+^) in the BM of *CT* mice treated with different regimens (n=6 per group). **g** qPCR analysis to detect the expression of *Elane* and *Il1b* in Mac1^+^ myeloid cells isolated by flow cytometry from the BM of mice treated with the different regimens (n=4). **h-i** Surface expression of Cd18 on granulocytes (**h**) and CCR2 on monocytes (**i**) isolated from different biological compartments was assessed by flow cytometry (n=4). **j** Histopathological analysis of colons collected from mice treated with different regimens (n=4-5 per groups). A score on a scale 1-4 was attributed by a trained veterinary pathologist to each individual mouse based on the infiltration of inflammatory cells in the colon (left) and mucosal ulceration of the colon (right). **k** heatmap for the expression of representative genes for different steps of hematopoiesis in HSPCs isolated from the pooled BM cells of 5 mice treated with each regimen. Red box: lymphoid signature. Green box: myeloid signature. Blue box: erythrocyte signature. **l** UMAP display of HSPC scRNA-Seq analysis. Cluster numbers are indicated, as well as lineage signature based on gene expression (M: myeloid; E: Erythrocytes; L: Lymphoid; MP: Myeloid Progenitors; Early: Earliest population). **m** *Csf2rb* (top) and Ezh2 (bottom) expression at the single cell level in HSPCs from mice treated with different regimens as identified by Seurat.

To characterize the consequences of single and dual inhibition on HSPC transcriptional profile at the single cell level, we performed scRNA-Seq of HSPC cells isolated from the BM of mice subjected to the different regimens. Dual inhibition in DSS-treated animals decreases the frequency of cells with a myeloid signature and restores lymphoid and erythrocyte signatures (**Fig. 7k-l; Fig. S8b-c**). In addition, individual transcriptional network visualization reveals that dual inhibition reduces the number of cells with high granulocytic transcription factor activity, while increasing the activity of transcription factors supporting monocytic (Irf8), erythroid (Ctcf, Fli1, Erg Ets1 and Gata2) and lymphoid (Ikzf1) lineages (**Fig. S8d-g**). Finally, *Csf2rb* and *Ezh2* expressions remain elevated in the context of dual inhibition (**Fig. 7m; Fig. S8d**). Combined with the unaltered cell cycle activity of HSPCs (**Fig. 7e**), this latest observation suggests that dual treatment represses stress myelopoiesis in the context of persistent E2f activity.

## DISCUSSION

Our data uncover an intrinsic mechanism that governs the hematopoietic output of HSPCs in the context of their sustained exit from quiescence. Specifically, we identify the simultaneous activation of a canonical cell cycle program with the activation of non-canonical target genes *Csf2rb* and *Suz12*/*Ezh2* by E2f as a coordinated mechanism to drive HSPC proliferation while enforcing their differentiation towards the myeloid lineage at the expense of alternative hematopoietic lineages such as erythropoiesis and lymphopoiesis.

Comprehensive scRNA-Seq analysis reveals several consequences of sustained HSPC exit from quiescence for hematopoiesis. While disruption of quiescence generates an alternative state for the most primitive cells within the LT-HSC compartment, their multipotency is barely affected by their proliferative status. Instead, our data indicate that the myeloid skewing decision occurs in the MPP compartment, with an expansion of MPP4 population, a rewiring of their cellular fate exclusively towards myelopoiesis and their projection to a GMP-like status. Of note, the myeloid-biased differentiation of TKO MPP4 involves limited changes in chromatin structure, suggesting that cell fate alteration in proliferative MPP4 in response to stress is very reactive. Downstream of MPP, the absence of MP expansion and their shortened trajectory suggest that differentiation is simply accelerated through the MP compartment to increase the myeloid output (**Fig. 2** and **Fig. 6**).

E2f has predominantly been studied for its critical role in the regulation of cell cycle target gene expression^19^, despite evidences suggesting that E2f is poised to exert additional non-cell cycle functions in a cellular context-dependent fashion. Our data establishes E2f as a central regulator of stress myelopoiesis because of its capacity to coordinate proliferation, the promotion of myelopoiesis via the enhancement of βcytokine signaling and the repression of alternative hematopoietic fates via the activation of core components of the repressive PRC2 complex. From an evolutionary standpoint, we propose that granting the coordination of cell cycle and cell fate decision to a unique factor ensures a fast and efficient response of hematopoietic progenitors to external stresses (such as the surge of Il3 and Gm-Csf systemic expression in inflammatory contexts) that require rapid myeloid cell production. However, pharmacological targeting of βcytokine signaling and Ezh2 activity in the TKO and the DSS model show that E2f canonical (proliferation) and non-canonical (differentiation) functions can be uncoupled, which enhances their relevance as therapeutic targets. A limitation of this pharmacological approach is that it generates a systemic inhibitory effect. Although Ezh2 expression is reduced in the guts of colitis patients compared to healthy controls^52^, this experimental limitation formally precludes the conclusion that the restoration of colon homeostasis (**Fig. 7j**) is solely the consequence of repressed stress myelopoiesis.

While we postulate that this cell-intrinsic mechanism is independent of the nature of the extrinsic stimuli (cytokine, growth factor, etc) that trigger it, we hypothesize that the role of the extrinsic inflammatory stimuli driving stress myelopoiesis is two-fold: *i)* serve as a proliferative cue to disrupt the Rb/E2f interaction (mandatory); *ii)* superpose additional intracellular signaling to the mechanism activated by E2f to fine tune the production of specific myeloid subpopulations (optional). This potential multi-layer mechanism would explain the variability of the myeloid outputs observed in different inflammatory contexts, based on the nature of the stress and its associated pro-inflammatory cytokines.

Review of the literature (as mentioned in the Results section) as well as our own data suggest that increased *Csf2rb* expression in HSPCs is common to many types of inflammatory contexts. Combined with the established regulation of *Ezh2* by E2f, we postulate that the dual mechanism reported here is a hallmark of inflammation. As such, we propose that the development of novel molecules to combine the targeted inhibition of βcytokine signaling and Ezh2 activity could represent an efficient therapeutic strategy to treat a spectrum of inflammatory conditions.

## MATERIAL AND METHODS

### Mice

*Rosa26-CreER^T2^ TKO* and CT mice (mixed 129/Bl6 background) were previously described^28^ and Cre expression was induced by intraperitoneal injection of 1mg of Tamoxifen® (Sigma) in corn oil for five consecutive days. *Csf2rb* deficient mice were obtain from Jax laboratory (Ref 005940, Bl6 background)). Wild-type Bl6 mice were used as controls for experiments involving *Csf2rb* deficient mice. Mice were housed in LIMR animal facility. Inflammation models were generated by daily injection of Il1β (general inflammation model) and LPS (systemic infection model) as well as addition of DSS 1-2% w/v (colitis model; 2% for Bl6 mice and 1% for 129/Bl6 mice) to drinking water. Il1β (Prepotech), LPS and DSS (Sigma) were reconstituted with PBS. All experiments were approved by LIMR IACUC (protocol #A20-348). All histological analysis were performed by the Comparative Pathology Core at the University of Pennsylvania School of Veterinary Medicine.

### Flow Cytometry and Transplantation

Colon were digested as previously described in our liver studies^53^ to generate single cell suspension. Cells from the colon, BM, spleen and blood were filtered, and red blood cells were lysed with the ACK buffer (NH4Cl/KHCO3). The remaining white blood cells were stained with a cocktail of lineage antibodies (Cd3, Cd4, Cd8, B220, Ter119, Mac1 and Gr1) as well as antibodies against Kit, Sca1, Flk2/Flt3, Cd48, Cd150, FcgR, Cd34, Ly6c, Ly6g, Rank and Cd115 (all purchased from eBiosciences) and βchain receptor (Miltenyi). Inflammatory status of granulocytes and monocytes was determined by assessing the surface expression of Cd18 and CCR2 (eBiosciences), respectively. The surfaceome assay was performed with the LegendScreen from Biolegend. For transplantation, two million unfractioned BM cells were collected from CT and Rosa26-CreER^T2^ TKO mice and retro-orbitally transplanted into lethally irradiated (8G) *Rag 1^-/-^*mice. Recipient mice were injected with Tamoxifen five weeks after transplantation.

### Colony Assay, OP cell assay, liquid culture, lentiviral transduction and cytospin analysis

100 CT or TKO HSPC subpopulations or 10,000 BM cells were plated in 1 ml of methylcellulose, either pre-supplemented with cytokines (M3434) or not (M3334) (Stem Cells Technologies). Cytokines were obtained from Peprotech and resuspended in PBS. Colonies were counted after 9 days in culture. Colony cells were resuspended, counted and spun for histology analysis. For liquid culture, 50-200K MPP/MP cells were plated in 24 well plates in DMEM media supplemented with 10% serum, 20ng/ml of Il3, Il6 and TPO as well as 100ng/ml of SCF. MacI surface expression was assessed by flow cytometry after 5 days in culture. 100 CT or TKO HSPC subpopulation cells were cultured on OP9 cells for 10 days in triplicates. Their lineage identity was determined by flow cytometry, using Mac1 (myeloid) and B220 (lymphoid) as markers. Three lentiviral vectors expressing hairpins against genes of interest (*Csf2rb*, *Zbtb7b*) as well as the RFP marker were purchased from Vectorbuilder and the two most efficient vectors were used for assays. Lentiviral particles were obtained by transfecting 23T cells with lentiviral plasmids together envelop plasmids of third generation. Cells were spun with lentiviral particles at 2000rpm for 1hr. After 72hrs in culture, successfully transduced cells were flow-isolated based on their expression of RFP. Cytospin were obtained by spinning 20,000 cells at 500rpm against glass slides. Cells were stained with Giemsa stain, as described previously^28^.

### Ex vivo cytokine stimulation, ChIP, IP-Western and gene expression analysis

HSPCs were flow-isolated from CT and TKO mice two weeks after tamoxifen treatment and serum-starved for two hours prior to stimulation with 10ng/ml of Il3 and GM-Csf (Peprotech) for 10 minutes. Cells were fixed, permeabilized and stained for anti p-Erk (Cell Signaling) prior to analysis on LSR Fortessa. For ChIP assay, primary BM cells were collected into 0.5% SDS lysis buffer at a density of 50 x 10^6^ cells/mL and crosslinked in 2% formaldehyde for 10 minutes. Lysates were sonicated on a BioLogics Model 3000 Ultrasonic Homogenizer for 30 seconds at 40% power on ice to produce chromatin fragments of approximately 400-700bp. Antibodies against E2f1 and E2f3 were purchased from Santa Cruz. For immunoprecipitation (IP) assay, cells were lysed with RIPA buffer and incubated with either antibodies against HA or Zbtb7b (SantaCruz), before pulldown with Protein-A coated agarose beads (Santa Cruz). After washing, beads were resuspended in loading blue prior to processing by immunoblotting and detection of Cebpα, Cebpβ, Cebpδ, Cebpε, Xbp1, Pu.1 and Irf8 (Cell Signaling). For RT-qPCR, purified hematopoietic populations were sorted in Trizol LS and the extracted RNA was purified with the RNeasy Mini kit (Qiagen), followed by DNAseI digestion to minimize the risk of genomic contamination. Reverse transcription was performed with the Protoscript First Strand cDNA Synthesis kit (New England Biolabs) and qPCR was performed in duplicate with SYBR Green PCR Master Mix (Life Technologies) on the Viia7 Real-Time qPCR system (Life Technologies). Data were normalized using *gapdh* as a reference gene. Primer sequences are available upon request.

### In silico ATAC-Seq and 10x genomics

#### Sample preparation

*ATAC-Seq*: ATAC-seq libraries were prepared as described^54^. In brief, 50,000 cells were isolated by flow cytometry and washed with cold 1x PBS. Cell pellet were resuspended in 50 µl of cold lysis buffer (10 mM Tris-HCl, pH 7.4, 10 mM NaCl, 3 mM MgCl_2_ and 0.1% (v/v) Igepal CA-630) and immediately applied to centrifugation at 1,600x*g*, 4 °C for 10 min. Nuclei pellet was resuspended in 50 µl of transposition reaction mix (1x Tagment DNA Buffer, 2.5 µl of Tagment DNA Enzyme 1 (Illumina)) and incubated for 30 min at 37 °C. Subsequent steps of the protocol were performed as previously described (Buenrostro et al., 2013). Libraries were purified using a Qiagen MinElute Gel Purification kit and the concentrations were measured using both Qubit and KAPA qPCR. 2100 Bioanalyzer was used to check the quality of libraries. Libraries were sequenced on the Illumina NextSeq 500, with 75-bp paired-end reads. 10x genomics: cells were sorted by flow cytometry and resuspended at a concentration of 1000 cells/ul. Cell suspension were processed by the Next Generation Sequencing Core at Penn Medicine.

#### Data processing

ATAC-seq: after trimming the adapters with attack (version 0.1.5 – https://atactk.readthedocs.io/en/latest/index.html), the raw reads were aligned to the mm9 genome using bow-tie-1.1.2 ((https://genomebiology.biomedcentral.com/articles/10.1186/gb-2009-10-3-r25) with the following flags: --chunkmbs2000—sam—best—strata-ml-X2000. We used MACS2^55^ for peak calling with a q cutoff of 0.05. Downstream analysis and visualization was done using HOMER^56^ and deepTools2^57^. scRNA-seq: to process the raw data, we used cellranger (v 2.0.0), which includes read alignment and generating a gene by cell count matrix. The R package Seurat (v 2.3.0)^58^ was utilized for filtering out cells with high mitochondrial signal, clustering and visualization. The same list of markers for different hematopoietic subpopulations were display in heatmaps through the manuscript (**TableS7**). These representative genes were arbitrarily selected based on their expression in the hematopoietic system, as identified in Gene Expression Commons (https://gexc.riken.jp). Seurat do not display expression for a gene in a heatmap when its expression is low/undetectable in all cells of the corresponding dataset. In addition, cluster numbers are attributed based on their frequency within datasets, with the most abundant cluster arbitrarily designed as Cluster 0 by Seurat. Furthermore, single-cell trajectories were identified and examined using Monocle 2 (v 2.10.1)^59^, Monocle 3 (v 3_0.2.0)^60^ and RNA Velocity^61^. Finally, we reconstructed gene regulatory networks using the SCENIC (v 1.1.2-2) package^35^.

### Statistical Analysis

The expression level of 5038 Ezh2 target genes, as identified by SCENIC, was interrogated in eight different mature hematopoietic populations (Gene Expression Commons^33^). We first centered gene expression, as provided by Gene Expression Commons, by adding 100 to each value. For each gene, we determined the average of its expression through the eight hematopoietic populations and calculated the relative fold expression in each population. Means, medians, standard deviations and interquartile ranges of the relative fold expression for the 5038 genes was calculated for each population. A two-sample t-test was performed to compare the mean value among populations, with the Granulocyte population as the reference. A violin plot was produced of the median and IQR of the eight populations. Analysis was conducted with Stata 18.0 (Statacorp, LLC) and Excel.

## ACKNOWLEDGMENTS

We thank Florin Tuluc and the members of the Flow Cytometry at CHOP as well as the CHOP Single Cell Core Biology, the Comparative Pathology Core at Penn Veterinary School of Medicine and Penn Next Gen Sequencing Core. P.V. is supported by internal funds at LIMR, Sharpe Struma and Women’s Board grants at Main Line Health as well as an ISAC grant from the NIDDK.

## AUTHOR CONTRIBUTIONS

AC, GO, EK, YN, LH and JT designed and performed experiments; KEH performed bioinformatic analysis; SK performed statistical analysis; KT generated and sequenced ATAC-Seq libraries; NS served as a consultant; PV designed, performed and analyzed experiments and supervised the project; PV wrote the manuscript.

---

**Figure S1.**
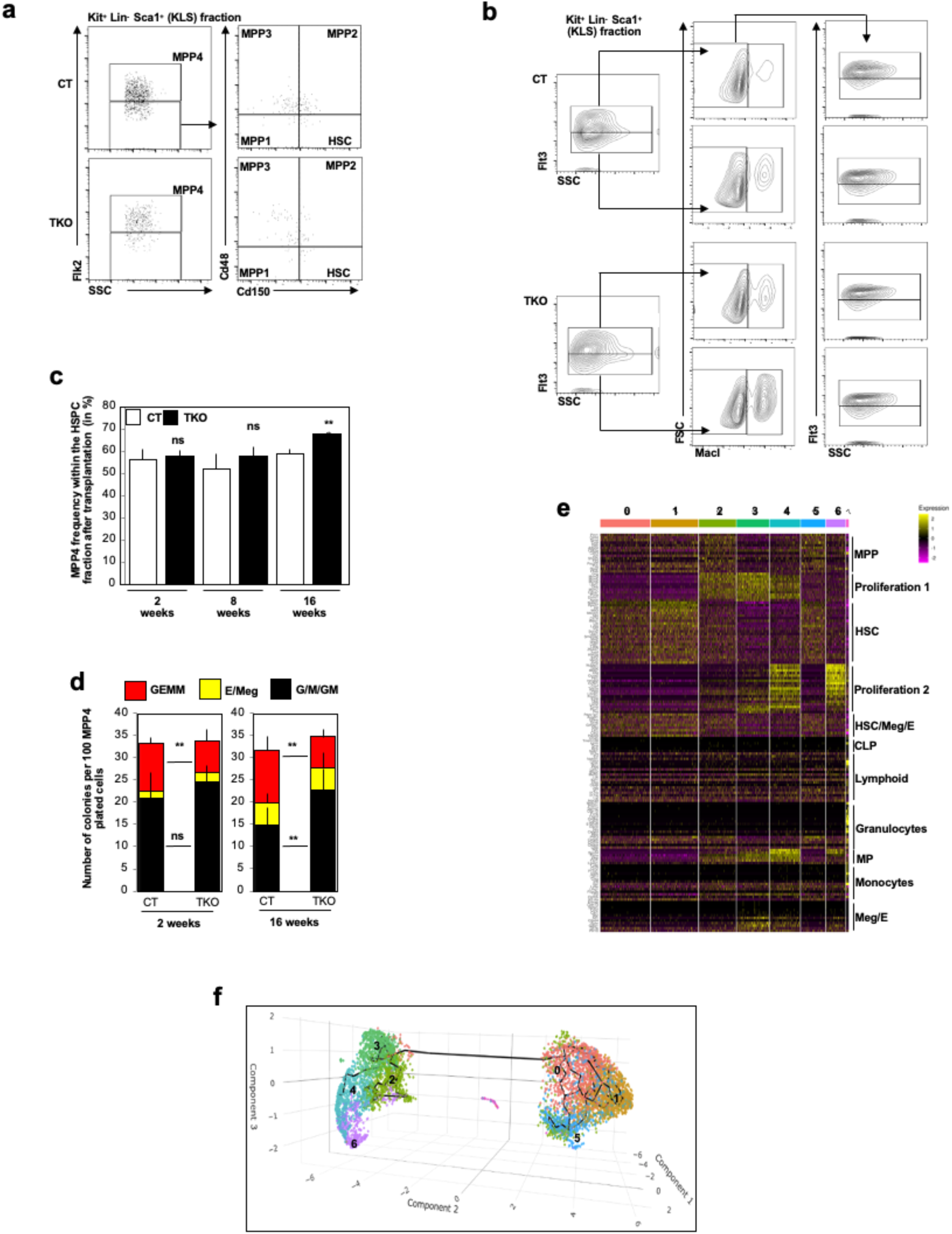
**a** Representative flow cytometry panel for Figure 1a**. b** Representative flow cytometry panel for Figure 1d. **c** Upon transplantation of *CT* and c*TKO* BM cells in *Rag1* deficient mice and subsequent Tamoxifen treatment upon the establishment of hematopoiesis (5 weeks), *CT* and *TKO* MPP4 frequency in the BM of recipient chimeric mice is evaluated at different time points (n=4). **d** 100 *CT* and *TKO* MPP4 isolated from chimeric mice at two- and 16-weeks post Tamoxifen treatment were plated into methylcellulose. Colonies were assessed after 8 days in culture (n=3). **e** Heatmap for selected genes expression for the merged *CT*/*TKO* LT-HSC dataset. **f** Trajectory analysis of merged *CT* and *TKO* LT-HSC datasets by Monocles-3D.

**Figure S2.**
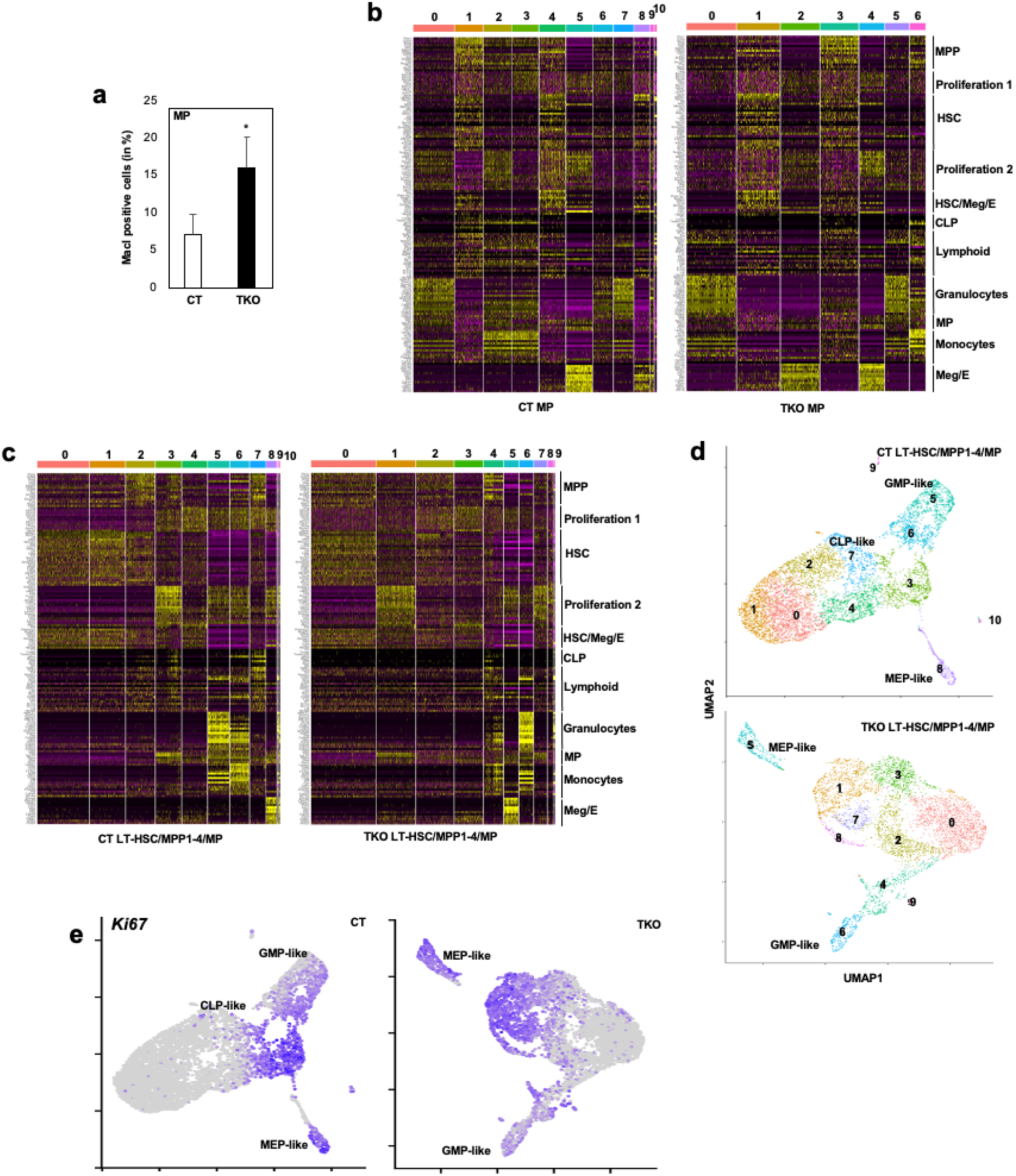
**a** *CT*/*TKO* MP were isolated by flow cytometry and placed in liquid culture. After 5 days, surface expression of the pan-myeloid marker MacI was assessed in cultured cells to quantify their differentiation rate. **b** Heatmap for the expression of representative genes for different steps of hematopoiesis in *CT* and *TKO* MP datasets. **c** Heatmap for the expression of representative genes for different steps of hematopoiesis in merged datasets for *CT* (left) and *TKO* (right) LT-HSC/MPP1-4/MP datasets. **d** UMAP display of merged CT (up) and TKO (down) LT-HSC-MPP1-4/MP datasets. **e** UMAP visualization of *Ki67* expression in merged *CT* (up) and *TKO* (down) LT-HSC/MPP1-4/MP datasets.

**Figure S3.**
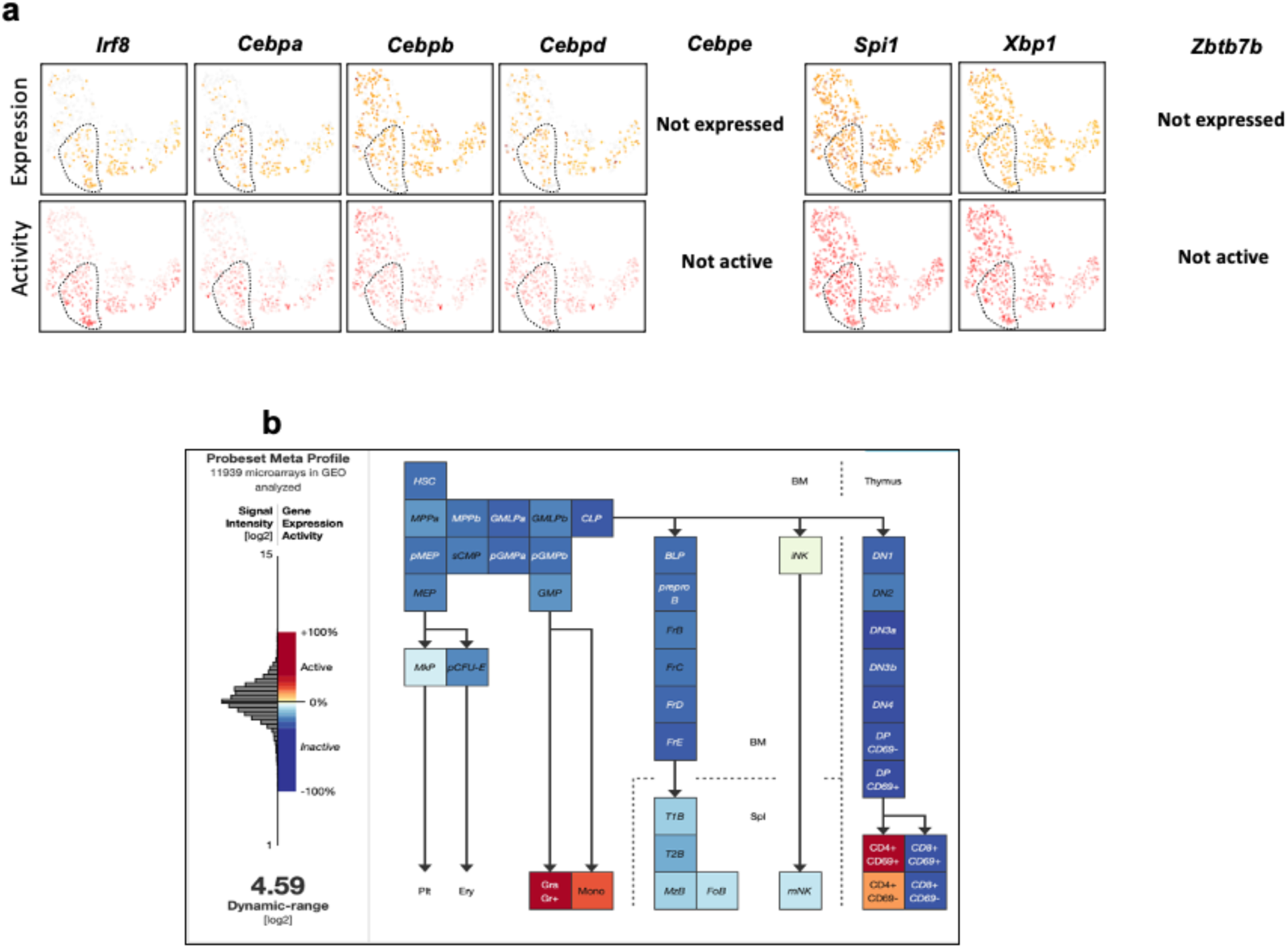
**a** Expression (up) and activity (down) level assessed by SCENIC for a myeloid network (*Irf8*, *Cebpa/b/d/e*, *Spi1* and *Xbp1*) as well as *Zbtb7b* in *CT* MPP4 are displayed at the single cell level. Cells corresponding to cluster 2 (lymphoid-primed) are circled with a black dotted line. **b** Gene Expression Commons analysis shows that *Zbtb7b* is exclusively expressed in mature myeloid and Cd4^+^ T cells in normal hematopoiesis.

**Figure S4.**
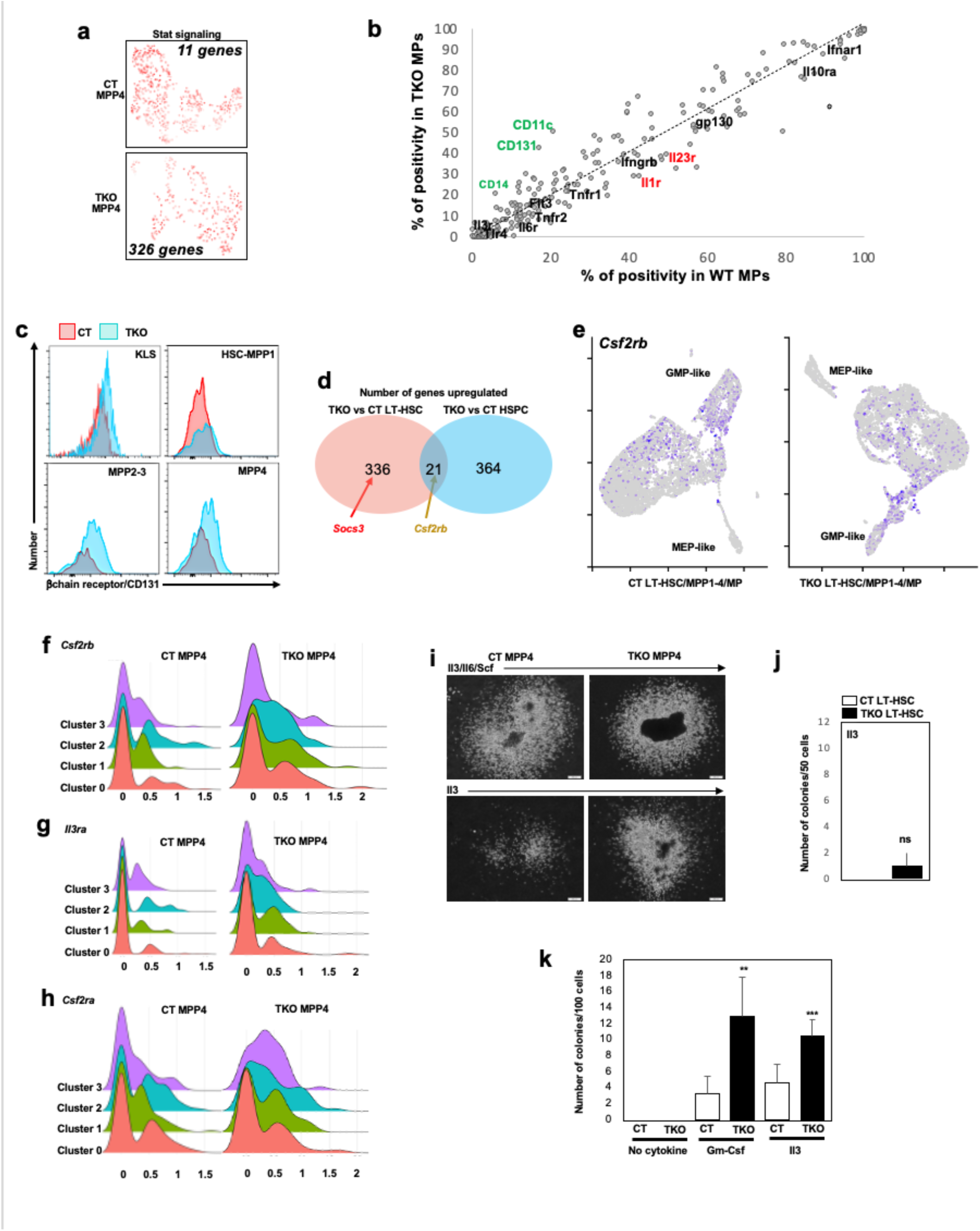
**a** *Stat* signaling transcriptional activity, as identified by SCENIC, in *CT* and *TKO* MPP4, including the respective number of *Stat* target genes (g) activated in each genotype. **b** Unfractioned BM cells from *CT*/*TKO* mice (5 per group) were isolated two weeks after Tamoxifen treatment, dyed for multiplexing purposes, merged, stained with antibodies against lineage markers (B220, Mac1, Gr1, Ter119, Cd3, Cd4, Cd8), Kit and Sca1 and screened for the expression of 256 surface markers (Biolegend antibody array® complemented by individual analysis for CD131 since it is not included in the array design). Background signal intensity was assessed by isotype staining. Each dot represents a surface marker expression in *CT* (x-axis) and *TKO* (y-axis) myeloid progenitors (MPs). The dotted line represents identical positivity % in *CT* and *TKO* MPs. Markers of monocyte lineage (CD14 and CD11c) and MPP4 (Flt3) are highlighted for technical validation. Gene names of interest are color-coded for better visualization when expression difference between *CT* and *TKO* HSPCs is at least 10 points. **c** Representative expression of the common βchain receptor (aka CD131, the protein product of *Csf2rb*) at the surface of *CT* and *TKO* LT-HSC/MPP1-4 subpopulations, as detected by flow analysis (n=3). **d** Venn diagram display of microarray expression analysis in *CT* and *TKO* progenitor populations: pink circle: LT-HSC; blue circle: HSPC cells. The figures indicate the respective number of genes upregulated in *TKO* vs *CT* cells. *Csf2rb* increased expression is observed in all *TKO* HSPC subpopulations while *Socs3* increased expression is restricted to *TKO* LT-HSC. **e** UMAP visualization of *Csf2rb* expression in merged *CT* (left) and *TKO* (right) LT-HSC/MPP1-4/MP cells. **f** Comparative expression of *Csf2rb* in each cluster from *CT* and *TKO* MPP4, as determined by scSeq-RNA analysis. **g-h** Comparative expression of *Il3ra* (**g**) and *Csf2ra* (**h**) in each cluster from *CT* and *TKO* MPP4, as determined by scSeq-RNA analysis. **i** Representative pictures of colonies from **Fig. 4e**. **j** 100 *CT* and *TKO* LT-HSC cells were plated in cytokine-free methylcellulose, supplemented with Il3 (n=3). Colonies were counted after 8 days in culture. **k** 100 *CT* and *TKO* MP cells were plated in cytokine-free methylcellulose, either not supplemented (no cytokine) or supplemented with Gm-Csf and Il3 (n=3). Colonies were counted after 8 days in culture.

**Figure S5.**
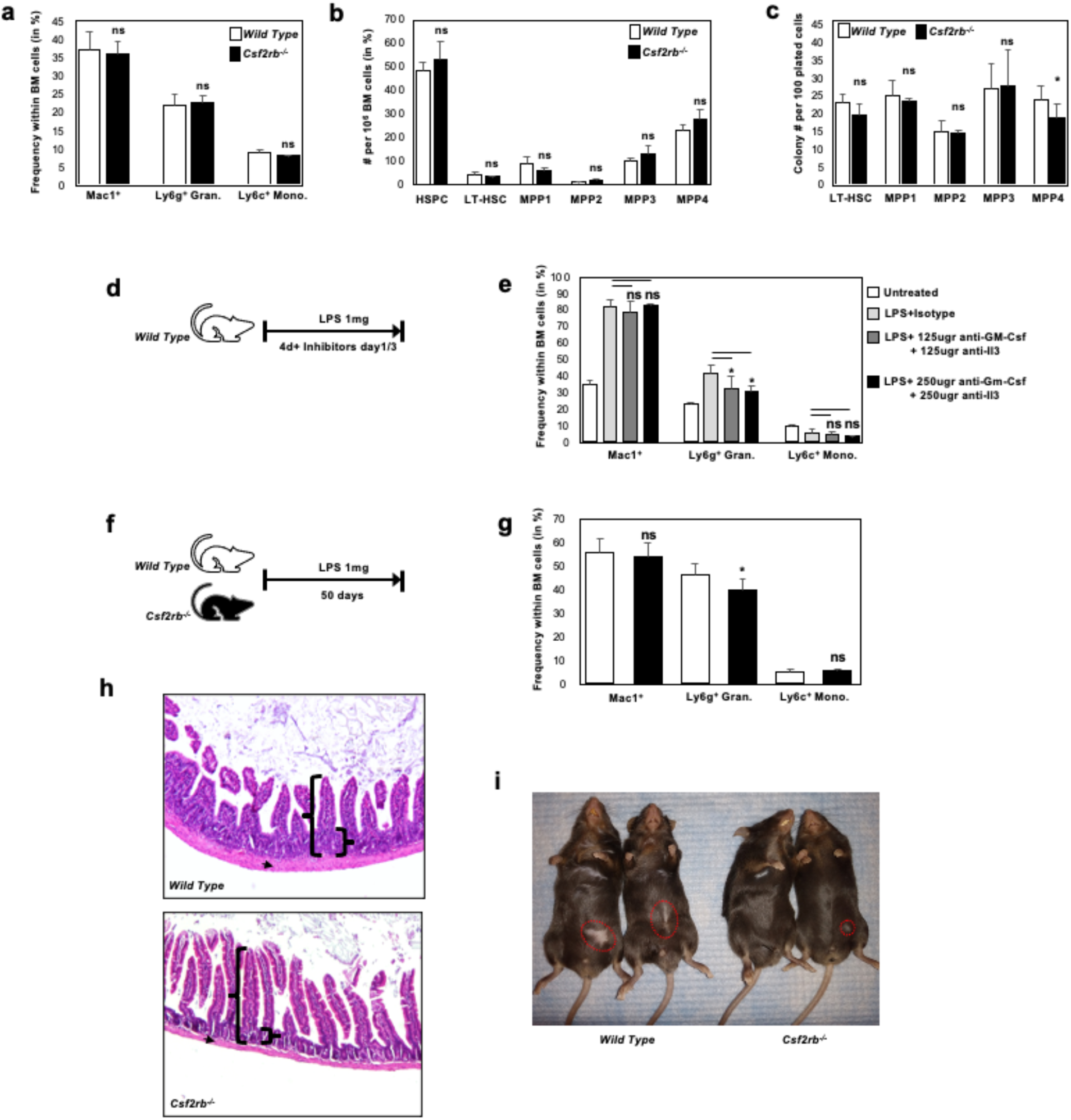
**a** Frequency of mature myeloid populations (Mac1^+^ myeloid cells, Ly6g^+^ granulocytes and Ly6c^+^ monocytes) in the BM of *wild type* and *Csf2rb* deficient mice (n=7). **b** Number of HSPC subpopulations per million BM cells in *wild type* and *Csf2rb* deficient mice (n=7). **c** Colony forming activity of HSPC subpopulations from *wild-type* and *Csf2rb* deficient mice in M3434® media. Colonies counted after 8 days in culture (n=3). **d** Wild-type mice were injected daily with LPS (1mg/kg) for 4 days to recapitulate an acute systemic infection. Mice were injected with different inhibitors on day 1 and 3 of the regimen. **e** Frequency of MacI^+^ cells, Ly6g^+^ granulocytes and Ly6c^+^ monocytes in the BM of mice from the different subgroups (n=4). **f** *Wild-type* and *Csf2rb* deficient mice were injected with LPS (1mg/kg) every day for 50 days to recapitulate a chronic systemic infection. **g** Frequency of MacI^+^ cells, Ly6g^+^ granulocytes and Ly6c^+^ monocytes in the BM of mice from the different subgroups (n=6). **h** Representative H&E staining of intestine sections from *wild-type* and *Csf2rb* deficient mice. Brackets: the ratio of villae/crypt is higher in *Csf2rb* deficient mice compared to *wild type*. Arrowhead: the basal membrane is thicker in *wild type* compared to *Csf2rb* deficient mice. **i** Representative pictures of scars (red dotted circles) developing at the site of injection in *wild type* and *Csf2rb* deficient mice.

**Figure S6.**
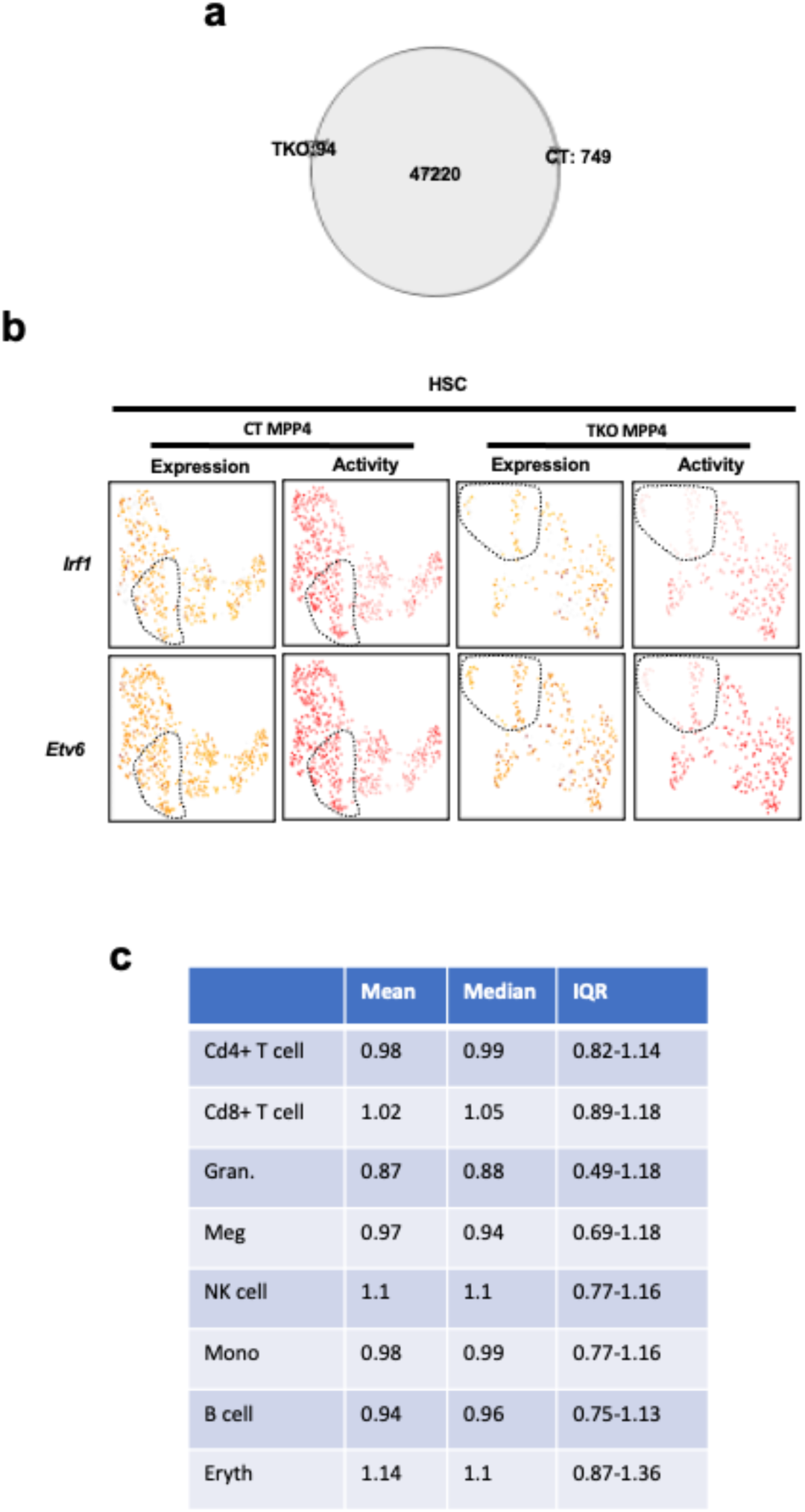
**a** Venn diagram displays the number of opened chromatin regions in *CT* and *TKO* MPP4, whether common to both conditions (47220) or unique to each condition (*CT*: 749; *TKO*: 94). **b** Expression and activity of representative transcriptional networks involved in the maintenance of stemness (HSC) as determined by SCENIC. **c** Statistical analysis of gene expression distribution for the 5038 Ezh2 target genes through 8 different mature hematopoietic populations, as determined by Gene Expression Common. IQR: Interquartile Range.

**Figure S7.**
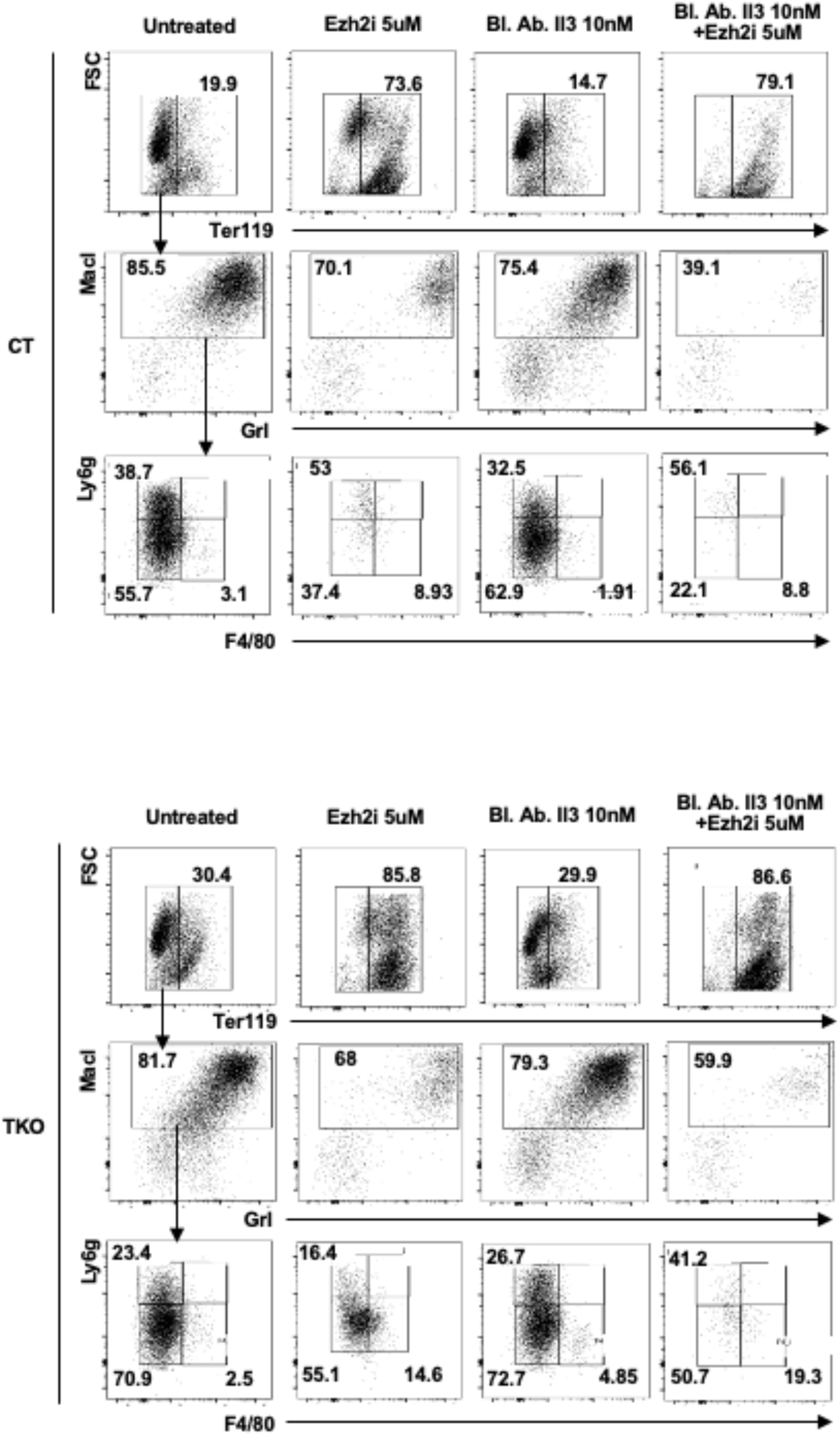
colony cells from selected *CT* and *TKO* conditions from **Fig. 6b** were resuspended and analyzed by flow cytometry for the expression of markers for myelo-erythroid populations (Ter119^+^: erythrocytes; MacI&GrI: myeloid markers; Ly6g^+^: granulocytes; F4/80^+^: monocytes). Numbers displayed are the average of three independent experiments.

**Figure S8.**
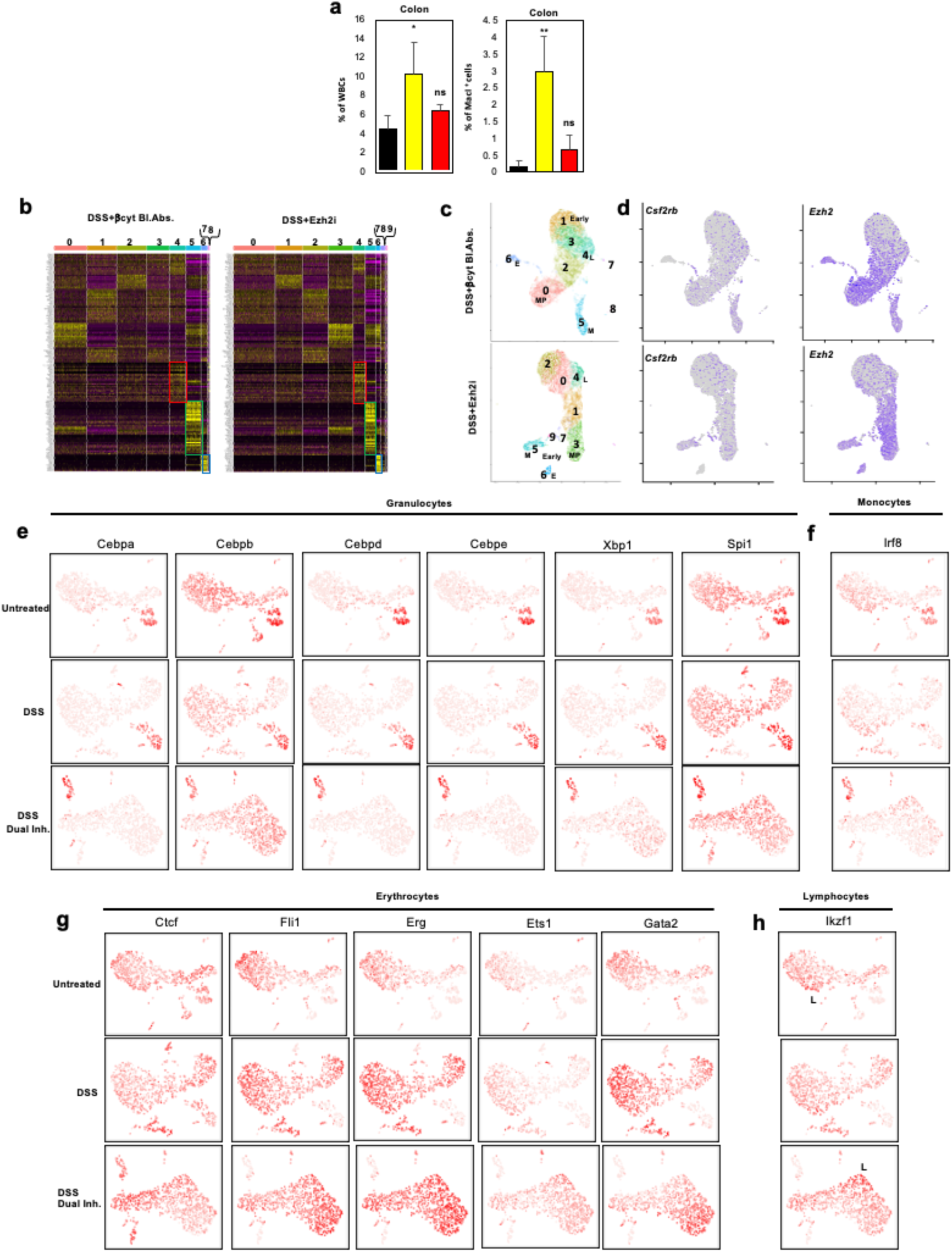
**a** Percentage of white blood cells (WBM-left)) and Mac1^+^ myeloid cells (right) in single cell suspension obtained by digestion of colon from mice exposed to different regimens (n=4). **b** heatmap for the expression of representative genes for different steps of hematopoiesis in HSPCs isolated from the pooled BM cells of 5 mice treated with each regimen. Red box: lymphoid signature. Green box: myeloid signature. Blue box: erythrocyte signature. **c** UMAP display of HSPC scRNA-Seq analysis. Cluster numbers are indicated, as well as lineage signature based on gene expression (M: myeloid; E: Erythrocytes; L: Lymphoid; MP: Myeloid Progenitors; Early: Earliest population). **d** *Csf2rb* (left) and Ezh2 (right) expression at the single cell level in HSPCs from mice treated with different regimens (DSS+βcyt inhibitors-up; DSS+Ezh2 inhibitor-down) as identified by Seurat. **e-f** SCENIC analysis displayed at the single cell level for granulocyte (**e**) and monocyte (**f**) transcription factors in mice either untreated, treated with DSS 1% or treated with DSS 1% and double inhibition for βcytokine signaling and Ezh2 activity. **g-h** SCENIC analysis displayed at the single cell level for erythroid transcription factors (**g)** as well as *Ikzf1* (*Ikaros*, a lymphoid transcription factor-**h**) in mice either untreated, treated with DSS 1% or treated with DSS 1% together with double inhibition for βcytokine signaling and Ezh2 activity. L: cluster associated with a lymphoid signature-see **Fig. 7j**.

**TableS1 genes expressed in clusters 0-3 of CT MPP4**

**TableS2 genes expressed in clusters 0-3 of TKO MPP4**

**TableS3 genes expressed in clusters 0-8 of merged CT MPP1-4& TKO MPP.**

**TableS4 genes expressed in clusters 0-9 of CT MP**

**TableS5 genes expressed in clusters 0-6 of TKO MP**

**TableS6 chromatin regions opened in CT and TKO MPP4**

**TableS7 markers used to identify hematopoietic subpopulations in heatmaps**

